# Endothelial PHD2 deficiency induces apoptosis resistance and inflammation via AKT activation and AIP1 loss independent of HIF2α

**DOI:** 10.1101/2024.02.01.578286

**Authors:** Shuibang Wang, Keytam S. Awad, Li-Yuan Chen, Mohammad A. H. Siddique, Gabriela A. Ferreyra, Caroline L. Wang, Thea Joseph, Zu-Xi Yu, Kazuyo Takeda, Cumhur Y. Demirkale, You-Yang Zhao, Jason M. Elinoff, Robert L. Danner

**Author notes:** To whom correspondence may be addressed: Critical Care Medicine Department, National Institutes of Health, 10 Center Drive, Room 2C145, Bethesda, MD 20892-1662, USA.

## Abstract

**BACKGROUND:** In hypoxic and pseudohypoxic rodent models of pulmonary arterial hypertension (PAH), hypoxia-inducible factor (HIF) inhibition reduces disease severity. However, HIF activation alone, due to genetic alterations or use of inhibitors of prolyl hydroxylase domain (PHD) enzymes, has not been definitively shown to cause PAH in humans, indicating the involvement of other mechanisms.

**METHODS:** Pseudohypoxia was investigated in primary human lung endothelial cells by silencing *PHD2,* and in *Tie2-Cre*/*Phd2* knockout mice, a rodent model of PAH. Lung vascular endothelial cells from PAH patients, and lung tissue from both SU5416/hypoxia PAH rats and PAH patients, were examined for validation.

**RESULTS:** *PHD2* silencing or inhibition, while activating HIF2α, induces apoptosis-resistance, hypo-proliferation, and IFN/STAT activation in endothelial cells, independent of HIF signaling. Mechanistically, PHD2 deficiency activates AKT and ERK, inhibits JNK, and reduces AIP1 (ASK1-interacting protein 1), all independent of HIF2α. Like PHD2, *AIP1* silencing affects these same kinase pathways and produces a similar dysfunctional endothelial cell phenotype, which can be partially reversed by AKT inhibition. These findings are corroborated in lung tissues of rodent PAH models and pulmonary vascular endothelial cells and tissues from PAH patients.

**CONCLUSIONS:** PHD2 deficiency in lung vascular endothelial cells induces an apoptosis-resistant, inflammatory, and hypo-proliferative phenotype. AKT activation and AIP1 loss, but not HIF signaling, drive these aberrant phenotypic changes. Our study suggests that HIF blockade alone may not suffice for PAH therapy; targeting PHD2, AKT, and AIP1 has the potential for developing more effective treatment.

**GRAPHIC ABSTRACT:** 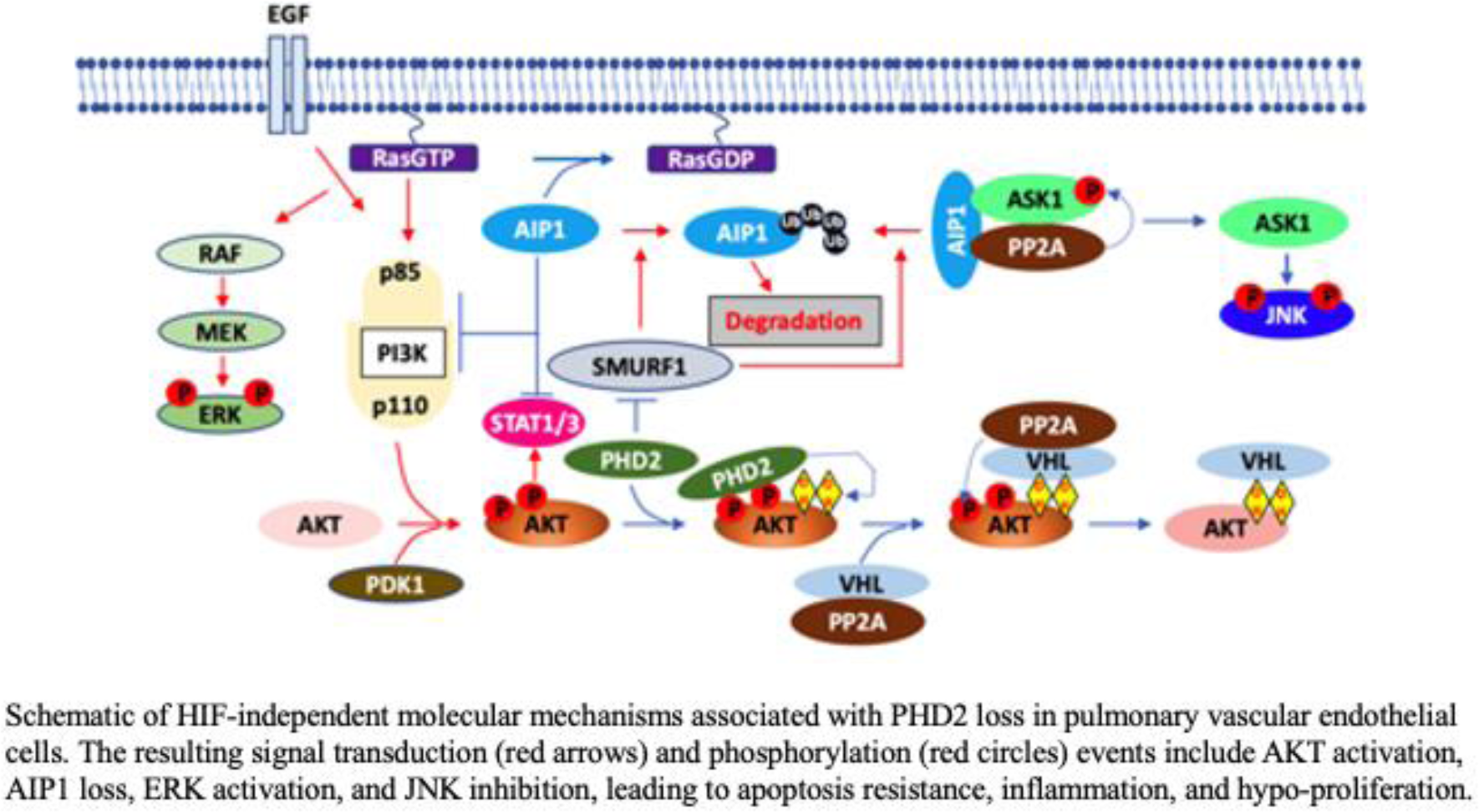

**Highlights:**

- *PHD2* silencing in human lung vascular endothelial cells suppresses apoptosis, inhibits proliferation, and activates STAT signaling, effects that persist despite HIF2α inhibition or knockdown.
- *PHD2* silencing activates AKT and ERK, inhibits JNK, and decreases AIP1, all independently of HIF2α
- Like PHD2, *AIP1* silencing led to similar alterations in kinase signaling and endothelial cell phenotypes, which are partially reversed by ATK inhibition.
- These *in vitro* findings align with observations in lung vascular endothelial cells and tissues from rodent models of PAH as well as PAH patients.

## 1 INTRODUCTION

Pulmonary arterial hypertension (PAH) is characterized by vascular cell proliferation, inflammation, and apoptosis resistance, leading to vessel wall thickening, luminal occlusion, and plexiform lesions.^1^ This pathologic remodeling progressively increases pulmonary vascular resistance that ultimately results in right ventricular failure. While vasodilator therapy has improved survival and quality of life, PAH patients continue to die of right heart failure or undergo lung transplantation.^1^ Thus, PAH therapeutics targeting molecular mechanisms driving vascular remodeling and pulmonary vessel loss are being sought.

In patients and animal models, a complex network of interacting pathways has been implicated in the pathobiology of PAH, making it difficult to distinguish primary or central targets from secondary or derivative processes. While hypoxia-inducible factor (HIF) signaling, triggered by hypoxia itself or simulation of a pseudohypoxic state have been studied, its importance in PAH pathogenesis and value as a mono-therapeutic target remains incompletely understood. Hypoxia is primarily sensed by prolyl hydroxylase domain (PHD) proteins, oxygen-dependent dioxygenases that hydroxylate HIF1α and HIF2α, targeting them for proteasomal degradation via von Hippel-Lindau (VHL)-mediated ubiquitination.^2, 3^ In mice, pulmonary hypertension (PH) induced by chronic hypoxia or a PAH-like phenotype caused by *Tie2-Cre*-mediated *Phd2 (Egln1*, official gene symbol) disruption are prevented by endothelial-specific deletion or heterozygous knockout of *Hif2α* (*Epas1,* official gene symbol).^4–7^ Likewise, a HIF2α gain-of-function mutation (G537R/W), which protects it from PHD2-mediated hydroxylation, causes autosomal dominant erythrocytosis, and PAH has been reported in 2 of 7 affected family members,^8^ while PH has been diagnosed in 2 additional cases.^9^ Further supporting the singular importance of HIF2α in the pulmonary vasculature, mice bearing this mutation also develop PH.^8, 10^ In addition, patients with autosomal recessive Chuvash polycythemia caused by an VHL inactivating mutation (R200W) are susceptible to PH but are also prone to pulmonary embolism,^11–13^ a separate potential contributor to elevated pulmonary pressures. Nonetheless, polycythemia and PH in murine models of Chuvash have been reversed by heterozygous deletion or therapeutic inhibition of HIF2α alone.^14, 15^

Although the above evidence strongly implicates HIF signaling and HIF2α specifically in the development of PH or PAH, other work suggests a more complex relationship between hypoxic/pseudohypoxic mechanisms and PAH that warrant further investigation. For example, HIF1α activation in pulmonary artery smooth muscle cells was conversely shown to lower vascular tone and protect mice against PH,^16^ and *Cdh5-Cre*-mediated *Hif2α* deletion, rather than overexpression, was found to promote PH.^17^ Furthermore, inherited PHD2 loss-of-function mutations, P317R, H374R, and D254H, caused erythrocytosis due to stabilized HIF2α in 6 affected patients but have not been reported to cause either PH or PAH.^18–20^ Similarly, 8 separate cases of erythrocytosis linked to novel hereditary heterozygous HIF2α, gain-of-function mutations, M535V/I, G537W, and D539E, did not present with PAH or PH.^21–23^ Moreover, daprodustat, an oral PHD inhibitor developed for the treatment of anemia in end-stage renal disease, stabilizes HIF1α and HIF2α, but so far, has not been associated with the development of PH.^24, 25^ Importantly, PHDs and VHL regulate other non-HIF targets, such as AKT^26^ and the B55α subunit of VHL-associated protein phosphatase 2A (PP2A).^27^ Proteins like AKT sit in consequential signaling networks that regulate proliferation, apoptosis, inflammation, cell migration and cytoskeletal architecture, processes highly relevant to pathologic vascular remodeling in PAH.^1, 28, 29^

Here, disruption of the PHD2/VHL pathway and the role of HIF2α was investigated in a pseudohypoxia model of endothelial dysfunction simulating aspects of PAH pathobiology. *PHD2* silencing produced an apoptosis-resistant, hypo-proliferative, and inflammatory cellular phenotype in both human lung microvascular (LMVECs) and pulmonary arterial endothelial cells (PAECs). HIF2α was stabilized and induced glycolytic gene expression but did not fully account for the phenotypic changes or aberrant signaling in these altered cells. AKT, which is regulated by PHD2/VHL independent of HIF2α^26^ and AIP1 (ASK1-interacting protein 1, official symbol *DAB2IP*), a Ras GTPase-activating (GAP) scaffolding protein that regulates a number of signaling pathways including AKT, ERK, JNK and STAT,^30^ were investigated to determine the mechanistic underpinnings of pseudohypoxic endothelial dysfunction. Finally, lung tissue from the SU5416/hypoxia (SuHx) rat and *Phd2 (Egln1)* knockout mouse models of PAH, as well as from patients with PAH were examined to further test the validity of our findings.

## MATERIAL AND METHODS

### Human primary lung endothelial cell culture

Human primary lung microvascular endothelial cells (LMVECs) and pulmonary artery endothelial cells (PAECs) from different donors were purchased from Lonza (Walkersville, MD. See Table S1). Cells were seeded onto plates and cultured in complete medium comprising of endothelial basal medium-2 (EBM-2) and supplement EGM^TM^-2 MV for LMVECs or EGM^TM^ for PAECs. The culture plates for PAECs were coated with type I collagen (50 μg/mL in 0.02 N acetic acid) prior to cell seeding. The basal medium EBM-2 and supplements were obtained from Lonza. Cells between passages 1-5 were used for experiments. LMVECs isolated from congenital associated PAH patients and failed donor (FD) were obtained from the Pulmonary Hypertension Breakthrough Initiative (PHBI) Penn Cell Center (Table S1).

For measurements of mitochondrial mass and superoxide of LMVECs, cells were plated at 0.8×10^5^ cells/well in 6-well plates and cultured for 16 h in complete medium (2 ml/well), followed by transfection with specific *PHD2* siRNA or scramble control (15 nM). Seventy-two hours post-transfection, cells were detached using 0.25% trypsin-EDTA (Thermo Fisher Scientific), washed and resuspended with PBS, and incubated for 30 min at *37*°*C* with 100 nM MitoTracker Green FM (Thermo Fisher Scientific) to stain mitochondria, and 3 μM MitoSOX Red (Thermo Fisher Scientific) to detect mitochondrial superoxide. Cells were then labeled with Live/Dead dye and analyzed by MACSQuant Analyzer 10 Flow Cytometer (Miltenyi Biotec, San Diego, CA).

### siRNA gene silencing and quantitative real-time PCR (qPCR)

*PHD2* and/or *AIP1*, *HIF2α*, *HIF1β*, *SMURF1* were knocked down in cells using 15 nM siRNA each, except for *SMURF1*, which was knocked down using 30 nM siRNA (ON-TARGET plus siRNA; Dharmacon, Lafayette, CO. See Table S2 for specific siRNA information). Control cells were transfected with non-targeting siRNA pool (ON-TARGET plus Non-targeting pool; Dharmacon). Transfection of siRNA into cells was performed by using DharmaFECT1 transfecting reagent (Dharmacon) as per the manufacturer’s protocol.

For qPCR, total RNA was extracted using RNeasy kits (Qiagen, Valencia, CA) and quantified by NanoDrop (BioLabNet, VA). cDNA synthesis was then performed using reverse transcription kits (Bio-Rad Laboratories, Hercules, CA). qPCR was carried out with TaqMan® probe/primers (Applied Biosystems; see Table S3 for Unique assay IDs). Beta-actin (*ACTB*), Beta-2 microglobulin (*B2M*), and ribosomal protein L13a (*RPL13A*) served as internal reference genes, and results were presented as geometric means ± SD.

### Western blotting and immunofluorescence (IF)

For Western blotting, whole cell lysates or rat lung homogenates were prepared on ice and protein concentrations were determined. Equal amounts of protein were loaded onto NuPAGE™ Bis-Tris Protein Gels (Thermo Fisher Scientific), following previously described procedures.^31^ For protein level normalization, Western blots were probed with a horseradish peroxidase conjugated antibody to beta-actin (Sigma, St. Louis, MO). Quantification of protein bands by densitometry was performed with Image Lab software, version 6.0.1 (Bio-Rad Laboratories).

For IF staining of human lung tissue sections, explanted lung tissue from IPAH patients and failed donor controls (FD) embedded in optimal cutting temperature compound were obtained from the Pulmonary Hypertension Breakthrough Initiative (PHBI, See Table S1 for donors’ information). Lung sections were air dried, fixed with 4% paraformaldehyde at room temperature for 7 min, blocked with 10% donkey serum/PBS and incubated with sheep anti-von Willebrand factor and rabbit anti-pAKT (S-473), anti-PHD2, or anti-AIP1 antibodies overnight at 4°C (see Table S1 for antibody information). Sections were then incubated with Alexa Fluor 488 donkey anti-sheep and Alexa Fluor 594 donkey anti-rabbit IgG (Jackson ImmunoResearch Laboratories; both at 1:200 dilution, see Table S1) at room temperature for 1 h. Nuclei were counter stained with Hoechst 33342 (Thermo Fisher Scientific; 1:5000) for 15 min at room temperature and mounted with Fluormount-G (SouthernBiotech, Birmingham, AL) for confocal microscopy. Fluorescence images were acquired using a SP8 DMI6000 confocal microscope (Leica Microsystems, Germany), equipped with 63x oil objective lens (numerical aperture of 1.4), and were analyzed in the Leica Application Suite X (Leica Microsystems).

### Cell apoptosis assay

Forty-eight hours post-transfection with specific siRNAs or scramble controls, cells were seeded at 5 ×10^3^ cells/well in 96 well plates and allowed to adhere for 6 h in complete medium (100 μl/well). The medium was then replaced with fresh complete medium or serum- and growth factor-free medium; specified inhibitors or vehicle control were added to the cell culture medium. Twenty-four hours later, caspase-3/7 activities in cells were measured using a luminometric assay kit Caspase-Glo 3/7 (Promega, Madison, WI). The amount of luminescence was measured on the Victor3™ multilabel reader (PerkinElmer Inc.).

Apoptosis was also evaluated by Annexin V / Propidium iodide (AV / PI) staining. Cells were plated at 0.8×10^5^ cells/well in 6-well plates and cultured for 16 h in complete medium (2 ml/well). Cells were transfected with specific siRNAs or scramble controls for 48 h. The medium was then replaced with fresh complete medium or medium without serum and growth factors; specified inhibitors or vehicle control were added to the cell culture medium. Twenty-four hours later, cells were detached by 0.25% trypsin-EDTA (Thermo Fisher Scientific), washed, and stained using the FITC-Annexin V Apoptosis Detection Kit (BD Pharmingen™, Franklin Lakers, NJ) as described in the manufacture’s protocol. Apoptotic cells (AV+ / PI-) were gated and counted using MACSQuant Analyzer 10 Flow Cytometer (Miltenyi Biotec).

### Cell proliferation assay

LMVECs and PAECs were plated in 96-well plates (4 ×10^3^ cells/well) and cultured for 16 h in complete medium (100 μL/well). The cells were then either incubated with different PHD2 inhibitors DMOG (100 μM), DFO (100 μM) or FG-4592 (50 μM) or transfected with specific siRNAs or scramble controls for 72 h. In experiments examining the effects of HIF2α inhibition, PT2567 (10 μM) or vehicle were added at 6 h post-transfection. Cell proliferation was evaluated by quantitating cellular DNA content with fluorescent CyQUANT^®^ Cell Proliferation Assay (Thermo Fisher Scientific, Waltham, MA). Cell proliferation was also evaluated by bromodeoxyuridine (BrdU) incorporation with chemiluminescent ELISA-BrdU assay (MilliporeSigma, Rockville, MD), in which cells were serum starved for 24 h starting at 48 h post-transfection or treatments and labelled with BrdU for 24 h in complete medium.

### Rodent models of PAH

The SU-5416/hypoxia PAH rat model (referred as SuHx rats) was established as previously described ^32^. Briefly, Sprague-Dawley rats (Charles River Laboratories; Wilmington, MA) were subcutaneously injected with SU5416 (25 mg/kg; Tocris; Minneapolis, MN), exposed to hypoxia (10% FiO2) for 3 weeks in a Biospherix Oxycycler (Biospherix Ltd.; Parish, NY) and then returned to normoxia for 7 weeks. At week 3 and week 8, lung tissue samples were collected and homogenized for Western blotting. All aspects of animal testing and care were approved by the Animal Care and Use Committee at the National Institutes of Health Clinical Center.

*Egln1^Tie^*^2^(*Phd2*CKO) PAH mice were established as previously described ^6^. Paraffin embedded lung tissue sections (5 μM thick) of the PAH mice and their wild type littermates (WT) were sent to Histoserv, Inc. (Germantown, MD) for immunohistochemistry (IHC) using anti-AIP1 as the first antibody, HRP-conjugated anti-rabbit as secondary antibody, and DAB (3,3′-Diaminobenzidine) as a chromogen substrate. Images were scanned and analyzed with NDP.view2 (Hamamatsu Corp., Bridgewater, NJ).

### Microarray experiments and analysis

Microarray analysis was conducted on samples from five independent experiments using LMVECs from the same donor at the same passage number to minimize variability. Cells were transfected with either scrambled pool non-targeting control siRNA (siCTRL), *PHD2* (siPHD2, 15 nM), *AIP1* (siAIP1, 15 nM), or both (siBoth) siRNA for 48 hours, and total RNA was then extracted using RNeasy miniprep kits (Qiagen). RNA quantity and quality were assessed using the NanoDrop 8000 Spectrophotometer (ThermoFisher Scientific) and RNA 6000 Nano LabChip (Agilent 2100 Bioanalyzer, Santa Clara, CA), respectively. All samples exhibited intact 18S and 28S ribosomal RNA bands (RIN 8.6-10, 260/280 ratios 2.00-2.04) and underwent microarray expression analysis using Human Clariom™ S Assay (ThermoFisher Scientific). Microarray data were RMA-normalized and log2-transformed in ThermoFisher Transcriptome Analysis Console, annotated with Human Clariom™ S Assay, and imported into R for PCA and outlier detection. Both raw and normalized data were submitted to GEO (GSE250030) and comply with MIAME. Limma fitted linear mixed models with fixed effects (siPHD2, siAIP1, their interaction) and a random effect (replication) to expression data for each gene. Differentially regulated genes were identified using a false discovery rate (FDR) of 0.01 and a 1.5-fold change cutoff for siPHD2 and siAIP1 main effects, and an FDR of 0.1 for interaction effects. This resulted in 1,728 unique differentially regulated genes, of which 853 were regulated by siPHD2, 681 by siAIP1, and 928 by interaction, with overlap among the groups. These groups were then upload for ingenuity pathway analysis (IPA^®^).

### Quantification and statistical analysis

Differences of relevant gene expression (mRNA and protein), protein phosphorylation, MitoSOX, and MitoTracker between control and PHD2 siRNA transfected human LMVECs or PAECs were analyzed by paired t-test. Differences between lung tissues or cells from control and Sugen rats or from PAH patients and non-PAH were examined by pooled t-test. Dose-dependent effects of DMOG on caspase3/7 activities, protein levels and protein phosphorylation were tested by fitting linear regression model with LMVEC or PAEC donors as random effects. ANOVA followed by post hoc t test was used to analyze effects of PHD2 silencing on cell apoptosis, protein phosphorylation, gene expression, and cytokines levels in the presence and absence of various siRNA and pathway inhibitors. Statistical analysis was performed using JMP version 16 (SAS Institute) with a two-tailed α of 0.05 as the cut-off for significance. Data are represented as arithmetic means ± SE except mRNA levels measured by qPCR, for which geometric means of FCs were plotted.

## RESULTS

### *PHD2* (*Egln1*) silencing induces glycolytic genes and apoptosis resistance in LMVECS

To determine the effect of PHD2 deficiency on HIF2α and HIF2α-regulated glycolytic genes in human LMVECs, *PHD2* was knocked down using a targeted siRNA approach (Figure 1A and B). As expected, HIF2α protein increased in *PHD2*-silenced LMVECs compared to control siRNA (siCTRL) transfected cells, while *HIF2α* mRNA was reduced (Figure 1A and B). Stabilization of HIF2α in *PHD2*-silenced LMVECs induced mRNA and increased protein expression of glycolytic genes *GLUT1*, *HK2*, *PKM2*, and *LDHA*. (Figure 1A and B). Consistent with previous work that demonstrated a decline in both mitochondrial reactive oxygen species and mitochondria DNA content in PHD2-deficient cells,^33, 34^ mitochondria-derived superoxide (MitoSOX) and mitochondrial mass (MitoTracker) were both decreased in *PHD2*-silenced LMVECs (Figure 1C), while the ratio of MitoSOX to MitoTracker was unaffected (Figure 1D).

**Figure 1.**
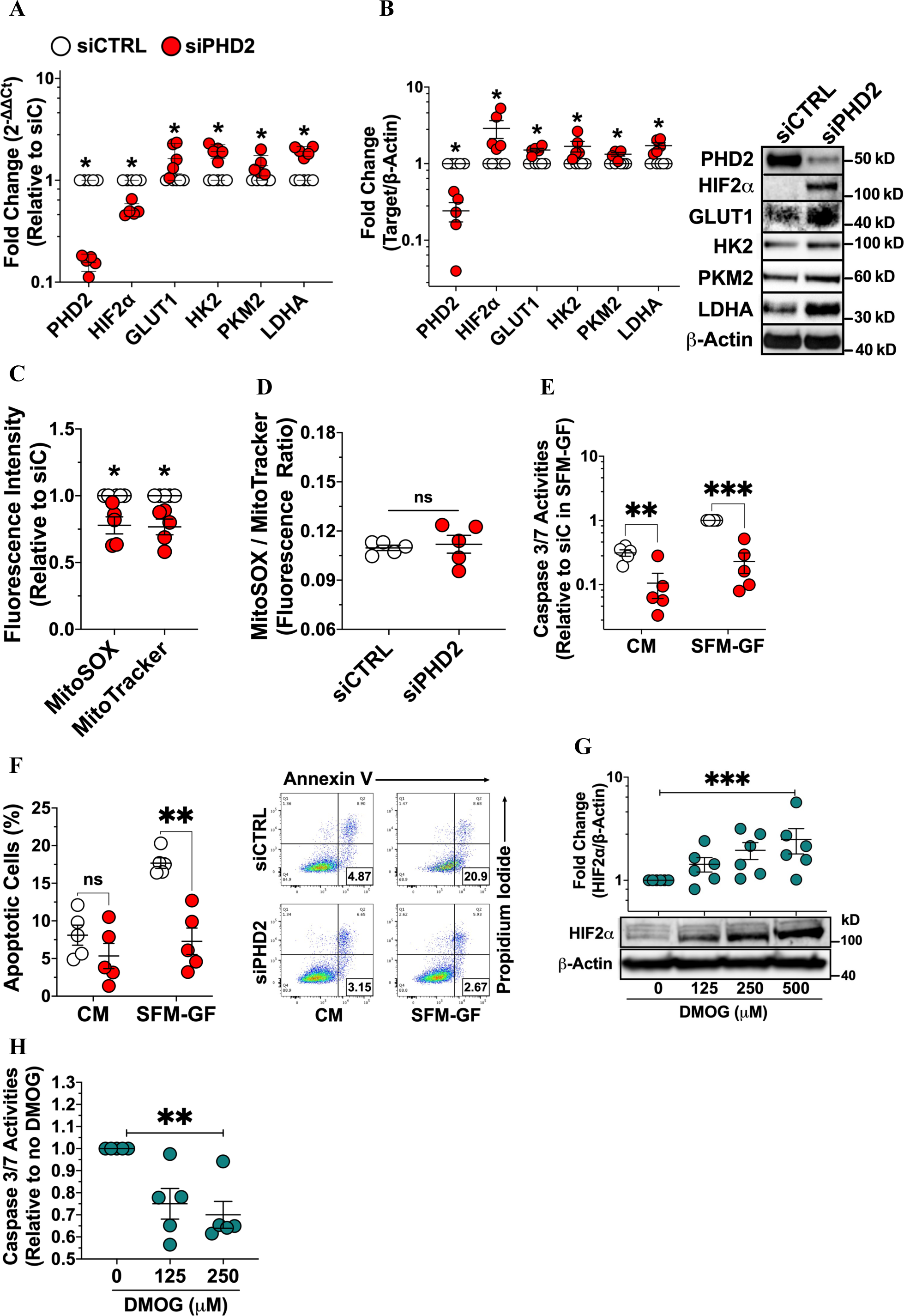
*PHD2* silencing induces glycolytic gene expression and apoptosis resistance in human lung microvascular endothelial cells. **A,** mRNA levels in cells transfected with control (siCTRL) or *PHD2* (siPHD2) siRNA for 48 h were measured by qPCR (n = 5). Data shown as geometric mean ± geometric SD. **B** through **D,** Protein levels were measured by Western blotting at 72 h post-transfection (n = 5). Mitochondrial superoxide and mass were analyzed by flow cytometry at the same time point (n = 5). **E** and **F**, Following 48 h transfection, cells were incubated in either complete medium (CM) or serum-free medium (SFM-GF) for 24 h. Apoptosis was assessed by caspase 3/7 assay and Annexin V/PI staining (n = 5). **G**, HIF2a protein levels were evaluated by Western blotting after 24 h treatment with DMOG, a PHD2 inhibitor (n = 6). **H**, Caspase 3/7 activity was measured in SFM-GF treated cells following 24h DMOG treatment (n = 5). Western blot protein levels were normalized to β-actin. Data are presented as mean ± SEM. *p < 0.05, **p ≤ 0.01, ***p ≤ 0.001. Paired t-test (A-D), two-way ANOVA with post-hoc t-test (E-F), linear regression (G-H).

Aberrant vascular remodeling in PAH has been associated with apoptosis-resistance, vascular cell proliferation, and reduced PHD2 expression.^6, 35, 36^ Here, decreased apoptosis in response to serum and growth factor withdrawal was confirmed in *PHD2*-silenced endothelial cells by caspase 3/7 assay (Figure 1E) and annexin-5/propidium iodine (AV/PI) staining (Figure 1F). Similarly, dimethyloxalylglycine (DMOG), a cell permeable PHD2 inhibitor, stabilized HIF2α protein and dose-dependently reduced caspase 3/7 activity (Figure 1G and H).

### Apoptosis resistance independent of HIF2α in *PHD2*-silenced LMVECs

Genetic deletion of *Hif2α* but not *Hif1α* reverses PAH and pulmonary vascular remodeling in *Tie2-Cre*-mediated *Phd2* knockout mice.^6, 37^ Therefore, PT2567, a specific HIF2α inhibitor, and HIF2α siRNA (siHIF2α) were used to investigate whether the aberrant phenotype of *PHD2*-silenced endothelial cells was entirely dependent on HIF2α. Surprisingly, PT2567 did not affect apoptosis in either *PHD2*-silenced LMVECs (Figure 2A) or *PHD2*-silenced PAECs (Figure S1A) as assessed by caspase 3/7 activation and AV/PI staining (Figure 2B), although it effectively blocked expression of known endothelial HIF2α targets GLUT1 and CXCR4 (Figure 2C). Unexpectedly, siRNA knockdown of *HIF2α*, which blocked the increase in HIF2α protein caused by PHD2 silencing (Figure 2D), did not prevent but rather enhanced apoptosis resistance, as evaluated by caspase 3/7 activation (p < 0.001 for the main effect of siHIF2α; Figure 2E) and AV/PI staining (p = 0.006 for the main effect of siHIF2α; Figure 2F) in LMVECs. No interaction was detected between *PHD2* and *HIF2α* silencing (p > 0.5 for the interaction; Figure 2E and F). Similar results were also found in PAECs (Figure S1B). Therefore, PHD2 silencing similarly induced apoptosis resistance in both the presence and absence of elevated HIF2α protein. To investigate whether HIF1α rather than HIF2α might have mediated apoptosis resistance induced by *PHD2* silencing, *HIF1β*, a constitutive co-factor for the HIFα family, was knocked down with *HIF1β* siRNA (Figure 2G). While *HIF1β* knockdown alone had no effect on caspase 3/7 activation (Figure 2H) and AV/PI staining (Figure 2I), co-silencing *HIF1β* and *PHD2* enhanced apoptosis resistance (p < 0.006 for interaction between *HIF1β* siRNA and *PHD2* siRNA; Figure 2H and I). Collectively, these results demonstrate that although *PHD2* silencing stabilizes HIF2α protein, the apoptosis-resistant phenotype associated with PHD2 deficiency is not mediated by HIF2α or other HIF family members in human LMVECs and PAECs.

**Figure 2.**
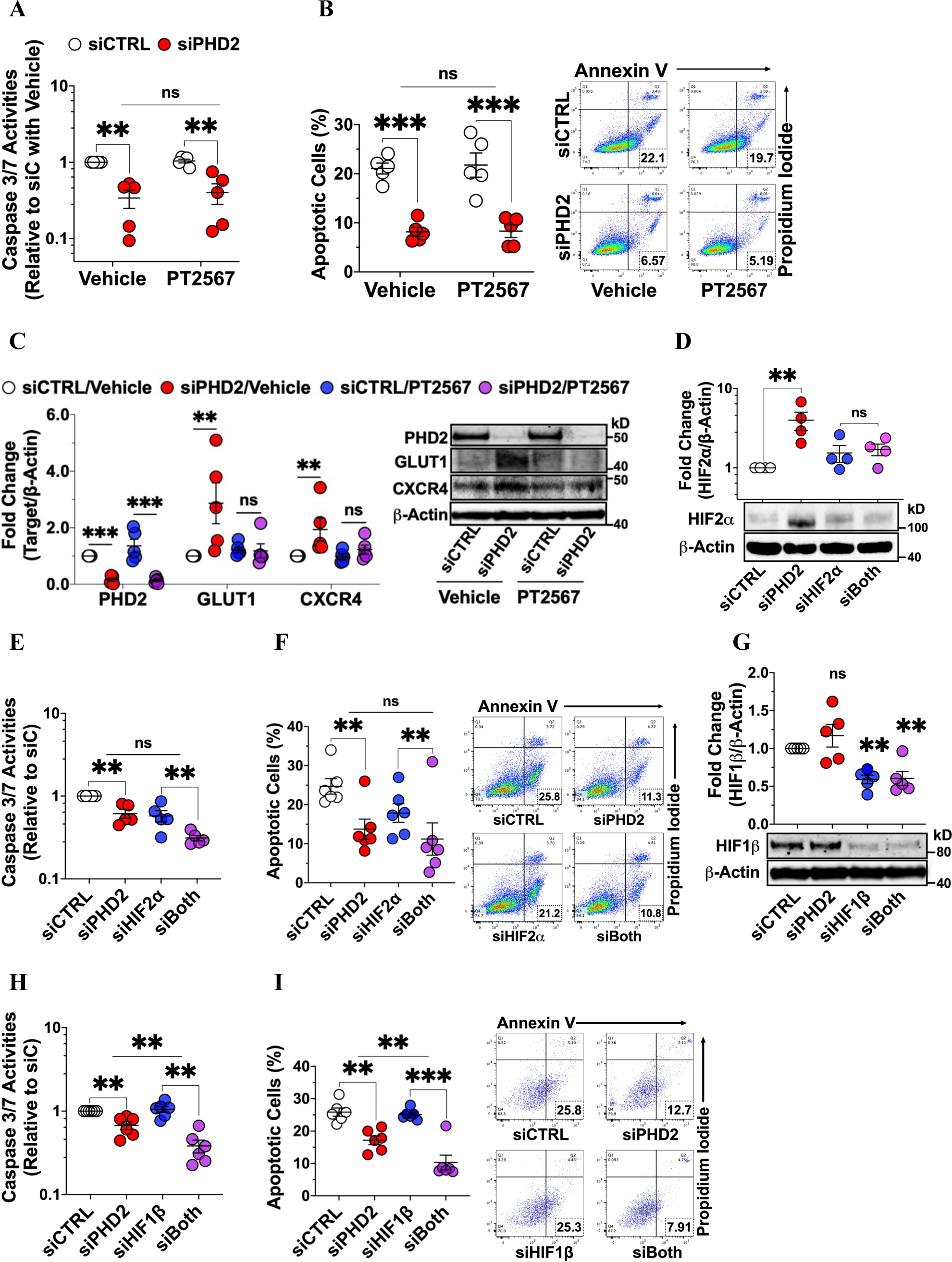
Neither HIF2α inhibition nor silencing reverses apoptosis resistance induced by *PHD2* silencing in human lung microvascular endothelial cells. Cells were transfected with siRNAs (siCTRL, siPHD2, siHIF2a, siHIF1b) or their combinations for 48 h **A** and **B,** Following 24 h incubation in serum-free media (SFM-GF) with PT2567 (HIF2a inhibitor) or vehicle, apoptosis was assessed by caspase 3/7 and Annexin V/PI staining (n = 5). **C**, Cells were further incubated for 24 h in complete media with PT2567 or vehicle for Western blotting (n = 5). **D**, Western blotting for cells 48 h post-transfection (n = 4). **E and F**, Following 24 h incubation in SFM-GF, apoptosis was assessed by caspase 3/7 and AV/PI staining (n = 5-6). **G,** Western blotting for cells 48 h post-transfection (n = 5). **H and I**, Cells were incubated and tested for apoptosis as in (E) and (F) (n = 5). Western blot protein levels were normalized to β-actin. Data are presented as mean ± SEM. **p ≤ 0.01, ***p ≤ 0.001. Two-way ANOVA with post-hoc t-test.

*Tie2-* or *Cdh5-Cre*-mediated *PHD2* knockout leads to PAH-like, proliferative, pulmonary vascular remodeling in mice.^6, 7^ In contrast, P*HD2* silencing or inhibition with DMOG, deferoxamine, or roxadustat (FG4592) in LMVECs and PAECs *in vitro* (Figure S2 and S3, respectively), produced a hypo-proliferative phenotype as assessed by quantifying cellular DNA content (CyQUANT^®^ Proliferation Assay) and BrdU incorporation. Moreover, the hypo-proliferative phenotype of *PHD2*-silenced LMVECs and PAECs (Figure S4 and S5, respectively) was not reversed by the HIF2α inhibitor, PT2567, or siHIF2α, both of which further reduced cell proliferation.

### *PHD2* silencing, independent of HIF2α, activates AKT and ERK, inhibits JNK, and activates STAT1/3 in LMVECS

Like HIF1α and HIF2α, phosphorylated (activated) AKT can also be directly hydroxylated by PHD2 and subsequently dephosphorylated (inactivated) by VHL-interacting PP2A.^26^ Thus, besides stabilizing HIF1/2α proteins, loss of PHD2 would be expected to activate AKT. AKT and ERK activation, as well as JNK inhibition, have been associated with apoptosis-resistance^29, 31, 38^ and/or IFN pathway activation.^29^ Therefore, we explored whether *PHD2* silencing affected these kinases in LMVECs. Compared to siCTRL transfected cells, AKT (pAKT-S473; Figure 3A) and ERK (pERK-T202/Y204; Figure 3B) were both activated, and JNK inhibited (pJNK-T183/Y185; Figure 3C) in *PHD2*-silenced LMVECs. Total AKT, ERK and JNK (Figure 3A through 3C) were not affected by *PHD2* silencing. Like *PHD2* knockdown, DMOG dose-dependently activated AKT and inhibited JNK (Figure S6A and S6B, respectively). In contrast to *PHD2* silencing, however, DMOG dose-dependently inhibited ERK activation (Figure S6C). Consistent with HIF2α independence, the specific HIF2α inhibitor, PT2567, did not alter AKT and ERK activation, or JNK inactivation in PHD2 deficient LMVECs (Figure 3D-3F, respectively).

**Figure 3.**
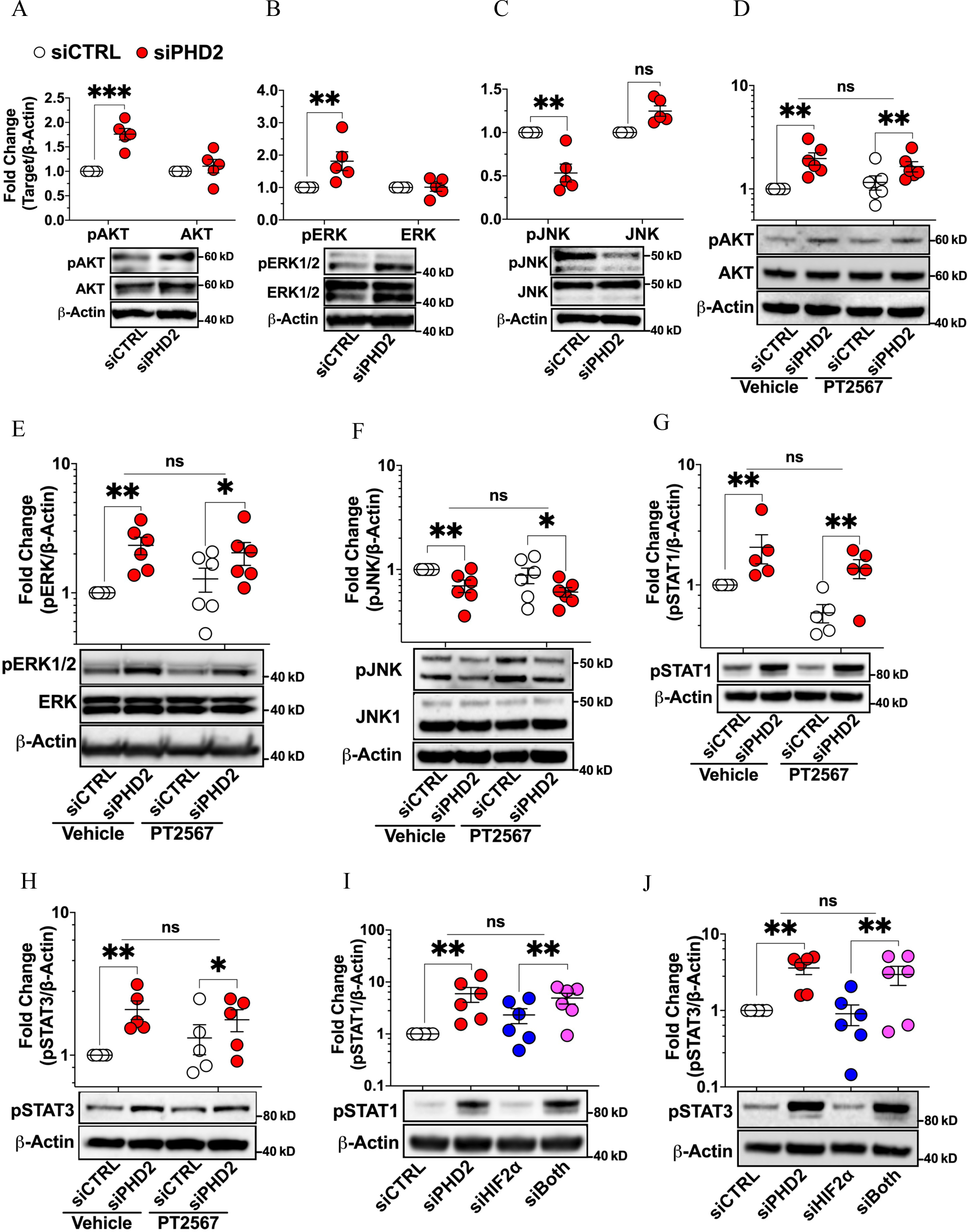
*PHD2* silencing, independent of HIF2a, activates AKT and ERK, inhibits JNK, and activates STAT1/3 in human lung microvascular endothelial cells. **A** through **C**, Cells were transfected with control (siCTRL) or *PHD2* (siPHD2) siRNA for 48 h. Activation of AKT (pAKT-S473), ERK (pERK-T202/Y204), and JNK (pJNK-T183/Y185) were assessed by Western blotting (n = 5). **D through H**, Cells were transfected as in (A) and incubated for another 24 h with HIF2a inhibitor PT2567 (10 mM) or vehicle control. Activation of AKT, ERK, JNK, STAT1 (pSTAT1-Y701), and STAT3 (pSTAT3-Y705 were assessed (n = 5-6). **I and J**, Cells were transfected with siCTRL, siPHD2, siHIF2a (HIF2a siRNA), or their combination for 48 h and incubated for another 24 h with PT2567 or vehicle. Activation of STAT1 and STAT3 were assessed (n = 6). Western blot protein levels were normalized to β-actin. Data are presented as mean ± SEM. **p ≤ 0.01, ***p ≤ 0.001. Paired t-test (A-C), two-way ANOVA with post-hoc t-test (D-J).

STAT1 and STAT3 have been implicated in the pathogenesis of PAH.^29, 36^ STAT1 stimulates inflammation, while STAT3 promotes cell survival and inhibits apoptosis. Hence, effects of *PHD2* silencing on STAT activation and its HIF2α dependence were also examined in LMVECs. Silencing *PHD2* activated both STAT1 (pSTAT1-Y701; Figure 3G) and STAT3 (pSTAT3-Y705; Figure 3H). Although PT2567 slightly inhibited STAT1 activation in both control and *PHD2*-silenced cells (P=0.005 for the main effect of PT2567; Figure 3G), neither the HIF2α inhibitor, PT2567 (Figure 3G and 3H), nor *HIF2α* silencing (Figure 3I and 3J) blocked the STAT1 or STAT3 phosphorylation that was induced by PHD2 loss.

### PHD2 loss reduces AIP1 protein expression and *AIP1* silencing mirrors the signaling and phenotype abnormalities observed in PHD2-deficient human LMVECS

AIP1 is a scaffolding Ras-GAP protein which inhibits both Ras effector pathways, namely Raf/ERK and PI3K/AKT.^30, 39^ Furthermore, AIP1 can activate JNK in endothelial cells by de-repressing ASK1 ^40^. Importantly, endothelial cell-specific *Aip1* knockout has been shown to cause vascular remodeling in mice.^41^ These previous findings and our present results led us to test whether *PHD2* silencing altered AIP1 expression. Compared to siCTRL transfected cells, AIP1 protein was markedly decreased by *PHD2* silencing (siPHD2) in LMVECs (Figure 4A) while AIP1 mRNA was not affected (Figure S6D), consistent with either reduced translation or increased degradation. Like *PHD2* silencing, DMOG dose-dependently decreased AIP1 protein (Figure S6E). The specific HIF2α inhibitor PT2567 did not alter AIP1 suppression in PHD2-deficient LMVECs (Figure S6F), demonstrating that the effect of PHD2 deficiency on AIP1 is independent of HIF2α. We next investigated the relevance of PHD2 loss-induced suppression of AIP1 to altered AKT, ERK and JNK activation in LMVECs. *AIP1* silencing (siAIP1) reduced AIP1 protein by 90% and did not affect PHD2 protein levels (Figure 4A). Like *PHD2* silencing, AIP1 knockdown activated AKT (p < 0.001 for siAIP1 *vs* siCTR; Figure 4B) and ERK (p = 0.002 for siAIP1 *vs* siCTR; Figure 4C), and suppressed JNK activity (p = 0.014 for siAIP1 *vs* siCTR; Figure 4D). Apoptosis resistance to serum and growth factor withdrawal was also detected in *AIP1*-silenced LMVECs (p < 0.001 and p = 0.007 for the main effect of siAIP1, respectively; Figure 4E and 4F). No interaction was observed between siAIP1 and siPHD2 on apoptosis (p > 0.5 for interaction in both; Figure 4E and 4F). AIP1 loss, like PHD2 loss, significantly inhibited human LMVEC proliferation (Figure S7A).

**Figure 4.**
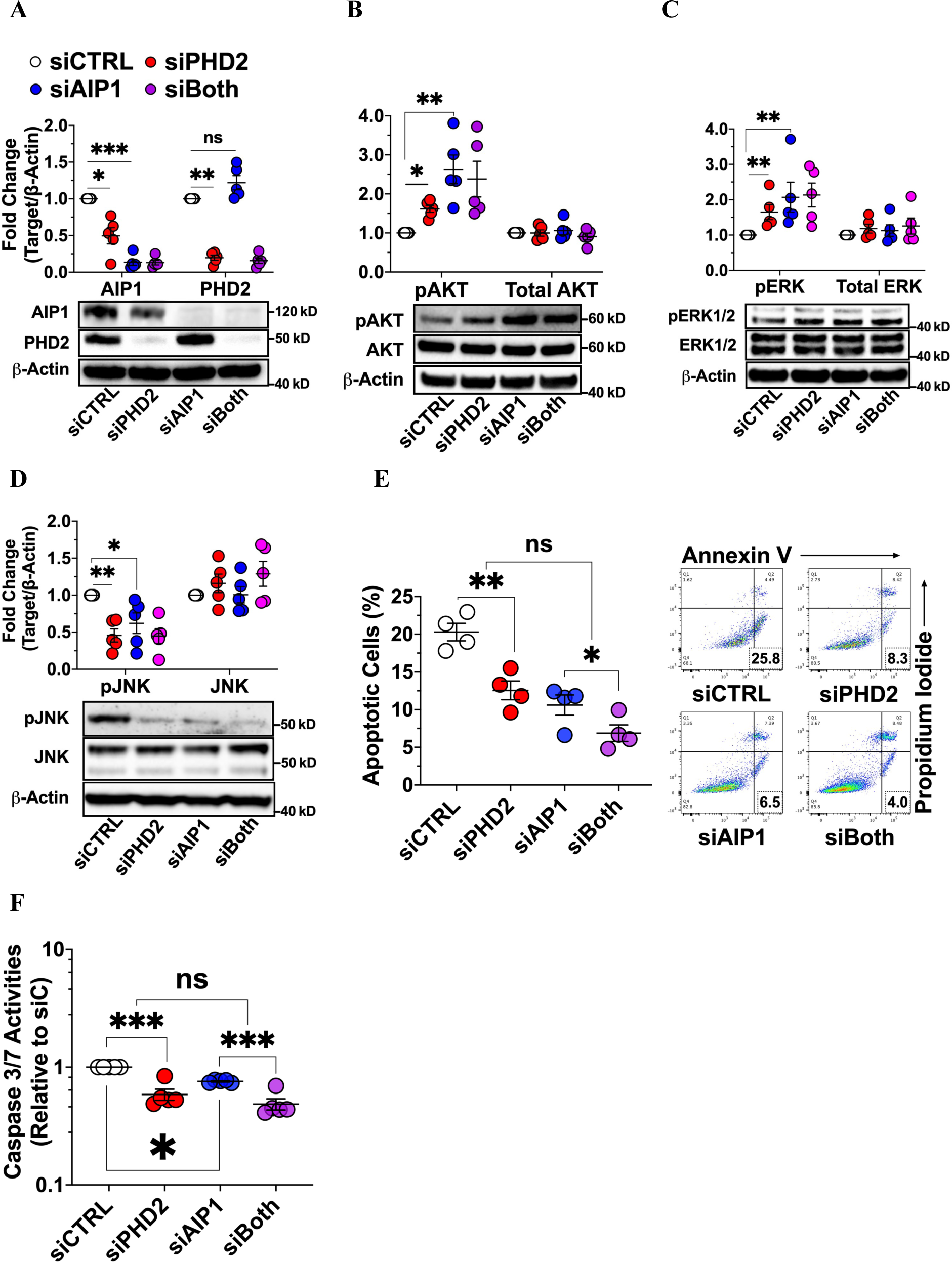
PHD2 loss reduces AIP1 protein expression and *AIP1* silencing activates AKT and ERK, inhibits JNK, and induces apoptosis resistance in human lung microvascular endothelial cells. **A** through **D**, Cells were transfected with control (siCTRL), *PHD2* (siPHD2), *AIP1* (siAIP1), or their combination for 48 h. AIP1 protein levels and activation of AKT (pAKT-S473), ERK (pERK-T202/Y204), and JNK (pJNK-T183/Y185) were assessed by Western blotting (n = 5). **E** and **F**, Cells were transfected as in (A), and incubated for an additional 24 h in serum- and growth factor-free medium and then assessed for apoptosis by Annexin V/Propidium Iodide staining and caspase-3/7 assay (n = 4-5). Western blot protein levels were normalized to β-actin. Data are presented as mean ± SEM. *p < 0.05, **p ≤ 0.01, ***p ≤ 0.001. Two-way ANOVA with post-hoc t-test.

### Loss of AIP1 activates STAT1/3, while inhibition of AKT reduces STAT1/3 activation and apoptosis resistance in *PHD2*- or *AIP1*-silenced human LMVECs

AIP1 has been shown to inhibit STAT1/3 in vascular smooth muscle cells (VSMCs), and STAT1 activation in endothelial cells has been ascribed to PI3K/AKT signaling.^29, 42^ Therefore, we hypothesized that, like PHD2, AIP1 deficiency in LMVECs might also activate IFN/STAT signaling. Silencing *AIP1* alone or in combination with *PHD2* knockdown, significantly activated STAT1/3 in LMVECs (Figure 5A). Leniolisib, a recently FDA-approved specific and well-tolerated PI3Kδ inhibitor,^43^ blocked STAT1/3 activation (Figure 5A). At the concentration used (5 μM), leniolisib inhibited AKT activation in both siCTRL transfected and *PHD*2-silenced LMVECs (Figure 5B). AKT inhibition promoted apoptosis as measured by AV/PI staining (p < 0.001 for leniolisib *vs* vehicle within siCTRL; Figure 5C) and caspase 3/7 activation (p < 0.001 for leniolisib *vs* vehicle within siCTRL; Figure S7B), and partly reversed apoptosis resistance induced by *PHD2* silencing (p < 0.001 for leniolisib *vs* vehicle within siPHD2; Figure 5C and S7B).

**Figure 5.**
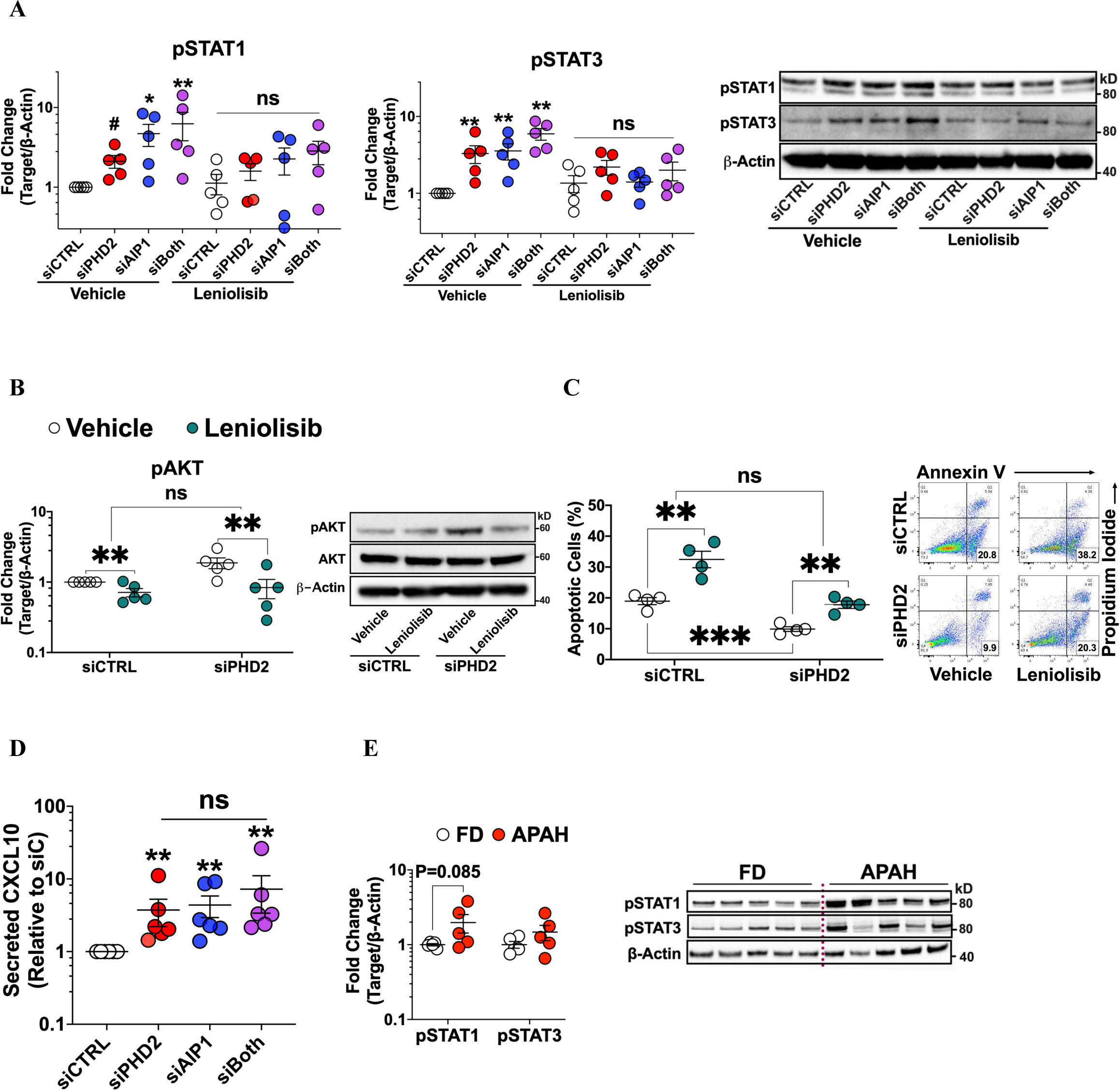
AKT inhibition inhibits STAT activation and reverses apoptosis resistance in *PHD2*- or *AIP1*-silenced human lung microvascular endothelial cells. Cells were transfected with scrambled control (siCTRL), *PHD2* (siPHD2), *AIP1* (siAIP1) siRNA or their combination. **A** and **B**, 48 h post-transfection, cells were further incubated for 24 h in complete medium with specific PI3Kd/AKT inhibitor leniolisib (5 mM) or vehicle control. Activation of STAT1 (pSTAT1-Y701), STAT3 (pSTAT3-Y705), and AKT (pAKT-S473) was assessed by Western blotting (n = 5). **C**, 48 h post-transfection, cells were further incubated for 24 h in serum- and growth factor-free medium with leniolisib or vehicle and assessed for apoptosis by Annexin V/Propidium Iodide staining (n = 4). **D**, 72 h post-transfection, cell culture supernatants were collected for measuring secreted CXCL10 levels by ELISA (n = 5). **E**, Lung microvascular endothelial cells isolated from congenital associated PAH (APAH) patients and failed donor (FD) controls were cultured in complete medium and assessed activation of pSTAT1-Y701 and pSTAT3-Y705 (n = 5). Western blot protein levels were normalized to β-actin. Data are presented as mean ± SEM. #p = 0.07; *p < 0.05; **p ≤ 0.01 Two-way ANOVA with post-hoc t-test (A-D), pooled t-test (E).

Consistent with STAT1/3 activation caused by PHD2 or/and AIP1 loss, secretion of CXCL10, an IFNγ signature gene, was induced by both *PHD2* and *AIP1* silencing in LMVECs (Figure 5D). Moreover, LMVECs isolated from congenital heart disease associated PAH patients (APAH) showed a trend toward increased STAT1/3 activation compared to control LMVECs from failed donor subjects (FD) (p = 0.085 for pSTAT1, p = 0.036 for the combined pSTAT1 and pSTAT3 effect; Figure 5E). These results suggest that loss of PHD2 and/or AIP1 in LMVECs leads to activation of the STAT/IFN pathway, and that signaling through AKT, independent of HIFα, contributes to this activation.

### Transcriptome analysis of human LMVECs after silencing *PHD2*, *AIP1*, or both

Transcriptomic profiling of LMVECs using microarrays identified 1,236 genes that were differentially regulated after *PHD2* (siPHD2, 853 genes) or *AIP1* (siAIP1, 681 genes) silencing [fold change (FC) ≥ 1.5 and false discovery rate (FDR) ≤ 0.01 for main effects; Figure 6A], and 928 genes that were regulated synergistically, either positively or negatively, by *PHD2* and *AIP*1 knockdown [FDR ≤ 0.1 for the interaction between siPHD2 and siAIP1 (siPHD2 X siAIP1); Figure 6A]. Although only 298 genes (298/1236 = 24.11%) were shared between siPHD2 and siAIP1 (Figure 6A), log2 fold change (LogFC) of 853 genes differentially regulated by siPHD2 and the LogFC of 681 genes differentially regulated by siAIP1 were positively and significantly correlated (r = 0.58, p < 0.0001; Figure 6B). The same was true for the LogFC between genes differentially regulated by either siPHD2 or siAIP1 and genes synergistically regulated by *PHD2* and *AIP1* silencing (Figure 6C and 6D). These results suggest that PHD2 loss and AIP1 loss have overlapping and concordant effects on gene expression in LMVECs through similar effects on specific signaling pathways.

**Figure 6.**
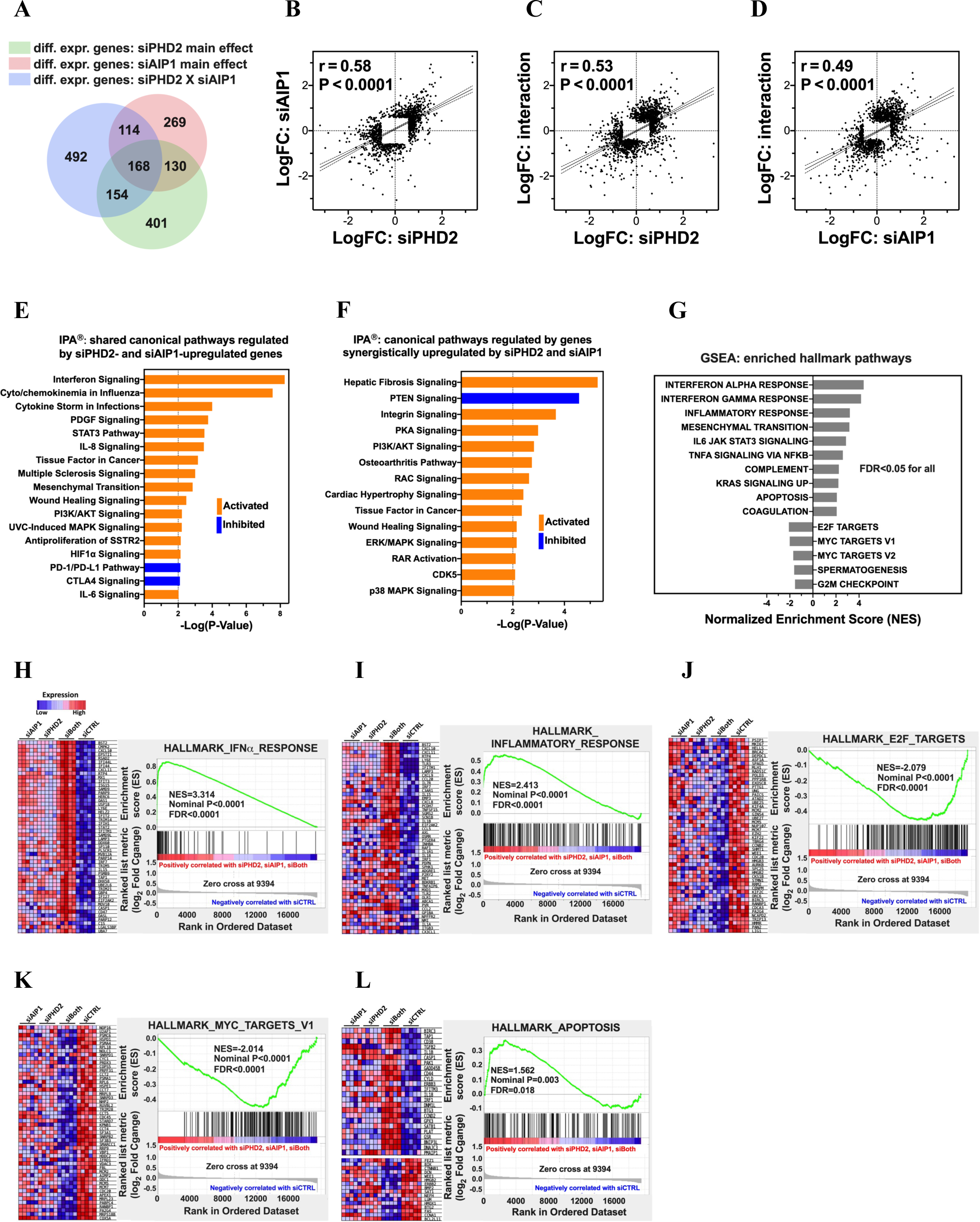
Transcriptome analysis of human lung microvascular endothelial cells silenced for *PHD2*, *AIP1*, and both using microarrays. **A**, Venn diagram of 1236 genes differentially regulated by silencing *PHD2* (siPHD2) or *AIP1* (siAIP1), 928 synergistically regulated, and their overlaps. **B**, Positive correlation between LogFC (Fold Change) of genes regulated by siPHD2 and siAIP1 (853 and 681 genes, respectively). **C** and **D**, Positive correlation between LogFC of synergistically regulated genes and genes regulated by siPHD2 or siAIP1. **E and F**, Top canonical pathways enriched by Ingenuity pathway analysis (IPA^®^, p ≤ 0.01) for genes upregulated by siPHD2 or siAIP1 (E) and genes synergistically upregulated (F). **G through L**, Gene set enrichment analysis (GSEA) revealed significant activation of IFNa and inflammation, and suppression of E2F and MYC targets by silencing of PHD2, AIP1, and both (siCTRL vs. siPHD2, siAIP1, and siBoth; FDR < 0.05). This was accompanied by the induction of anti-apoptotic genes (*BIRC3, TAP1, PAK1*) and repression of pro-apoptotic genes (*BCL2L11, FAS, BTG2*).

Ingenuity pathway analysis (IPA^®^) was used to identify canonical pathways associated with silencing of *PHD2*, *AIP1*, or both, by uploading our selected main effect lists of differentially up-(Figure 6E) and down-regulated (Figure S8A) genes. Multiple overlapping cellular signaling pathways were found to be significantly activated (Figure 6E) or inhibited (Figure S8A) by both *PHD2* and *AIP1* silencing. Among the significantly activated canonical pathways, IFN, cytokine/chemokine, STAT3, PI3K/AKT, and MAPK signaling (Figure 6E), IFN and cytokine/chemokine signaling had the most significant P values (Figure 6E). Similarly, for genes that were synergistically up-(Figure 6F) and down-regulated (Figure S8B) by *PHD2* and *AIP1* silencing, IPA^®^ identified PI3K/AKT and ERK/MAPK as significantly activated canonical pathways and PTEN signaling as inhibited (Figure 6F). Together, these IPA^®^ results support our experimental findings that loss of PHD2 or AIP1 in human LMVECs leads to activation of AKT, ERK and IFN/STAT.

Likewise, gene set enrichment analysis (GSEA), another pathway enrichment method that analyzes all genes without arbitrary selection cutoffs, also revealed that gene sets regulating IFNα/γ, inflammation, STAT3, KRAS, and apoptosis signaling were significantly enriched by silencing *PHD2* (siPHD2), *AIP1* (siAIP1), or both (siBoth) compared to control siRNA (siCTRL) (Figure 6G). In contrast, targets of E2F and MYC, well-known regulators of cell proliferation and apoptosis,^44, 45^ and the G2M checkpoint were significantly enriched in siCTRL (Figure 6G). Expression heat maps of the top 50 marker genes and enrichment plots of these pathways exhibited the activation of IFNα/γ (Figure 6H and Figure S8C), inflammation (Figure 6I), and KRAS (Figure S8D), and the suppression of E2F (Figure 6J) and MYC targets (Figure 6K), and the G2M checkpoint (Figure S8E) by silencing *PHD2* or *AIP1*, or both. As is visible in the heat maps, co-silencing had the strongest effects on these pathways. Among the enriched apoptosis-related gene sets, silencing of *PHD*2, *AIP1*, or both induced anti-apoptotic genes such as *BIRC3* (*C-IAP2*), *TAP1* (*PSF1*), and *PAK1*, while repressing pro-apoptotic genes including *BCL2L11*, *FAS, and BTG2* (Figure 6L). Together, these results demonstrated that PHD2 loss induced AKT activation and suppressed expression of AIP1 protein, with both effects contributing to an apoptosis-resistant, inflammatory, and hypo-proliferative pulmonary vascular endothelial phenotype.

### PHD2 loss and AIP1 dysregulation associated with AKT and STAT3 activation in rodent models of PAH

*PHD2* silencing in LMVECs reduced AIP1, activating AKT and STAT, which contributed to a PAH-like cellular phenotype. Consistent with these *in vitro* findings, PHD2 protein levels were reduced (Figure 7A), and both phosphorylated (Y705) and total STAT3 levels were increased (Figure 7B) in lung tissue homogenates obtained from SuHx rats 3 weeks after PAH model induction. At 3 and 8 weeks following model induction, a trend toward increased pAKT (S473) was observed in lung homogenates from SuHx rats relative to normoxia control rats (p = 0.09; Figure 7C) whereas total AKT expression was similar. In contrast to decreased AIP1 protein levels in *PHD2*-silenced LMVECs, whole lung AIP1 expression was increased in SuHx rats (Figure 7A). However, immunohistochemical staining of lung sections showed that AIP1 (brown staining) in *Egln1^Tie^*^2^(*Phd2* CKO) PAH mice was reduced compared to control wild type (WT) mice (Figure 7D). Differences (red arrowheads) in AIP1 staining between WT and PAH mice were seen within the endothelial lining of small pulmonary blood vessels (Figure 7D).

**Figure 7.**
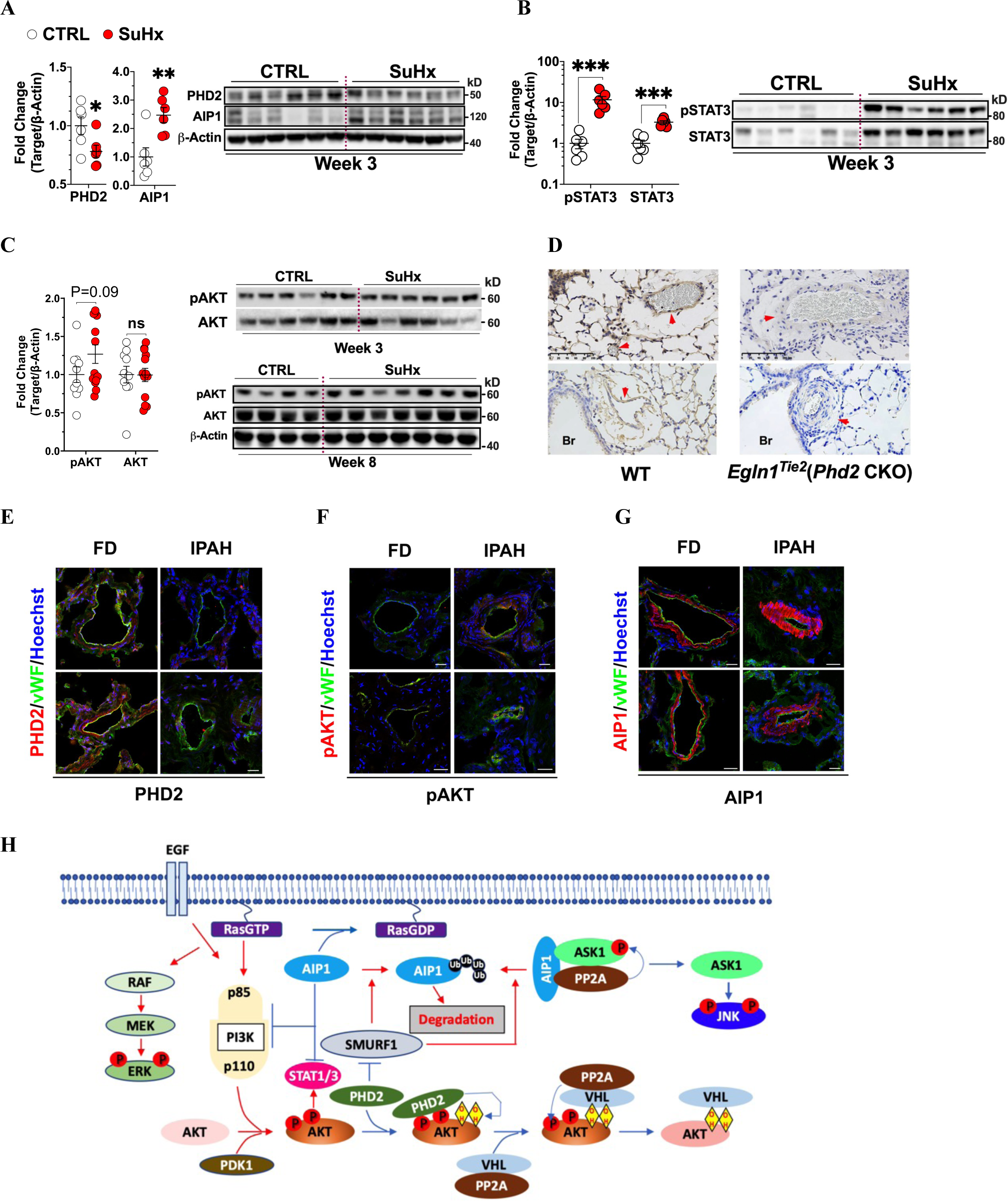
Diminished PHD2, dysregulated AIP1, and activated STAT and AKT in lungs of pulmonary arterial hypertension (PAH) animal models and in lung endothelial cells of PAH patients. **A** through **C,** PAH induced in SD rats (SuHx) by SU5416 followed by 3 weeks of hypoxia and then 5 weeks of normoxia. Lung protein levels of PHD2, AIP1, pSTAT3 (Y705), STAT3, pAKT (S473), and AKT were measured at week 3 and 8 vs. normoxic controls (CTRL). Western blot analyses were normalized to β-actin (mean ± SEM). **p ≤ 0.01, ***p ≤ 0.001. Pooled t-test (n = 6-13). **D**, Mice with conditional *Phd2* knockout [*Egln1^Tie^*^2^(*Phd2* CKO)] developed PAH. Immunohistochemistry revealed reduced AIP1 expression in the lungs of PAH vs. wild-type controls (WT). Representative images from four independent experiments are shown and vessels (red arrowheads) and bronchioles (Br) indicated. Scale bar, 100 μm. **E** through **G**, Lung sections from idiopathic PAH (IPAH) patients and failed donor (FD) control were co-stained for von Willebrand factor (vWF, green), PHD2, pAKT (S473) or AIP1 (red), and counterstained with Hoechst 33342 (blue). Representative immunofluorescence images from five different IPAH patients and FD Controls are shown. Yellow staining represents co-localization of PHD2, pAKT and AIP1 in lung vascular endothelial cells. Scale bar, 25 μm. **H**, Schematic of HIF-independent mechanisms associated with PHD2 loss in pulmonary vascular endothelial cells. The resulting signal transduction (red arrows) and phosphorylation (red circles) events include AKT activation, AIP1 loss, ERK activation, and JNK inhibition, leading to apoptosis resistance, inflammation, and hypo-proliferation

### Diminished PHD2 and dysregulated AIP1 associated with AKT activation in the pulmonary endothelium of patients with PAH

Immunofluorescent co-staining for PHD2 or pAKT (S473) and von Willebrand factor (vWF), an endothelial cell marker, demonstrated that PHD2 was diminished (Figure 7E), while pAKT was increased (Figure 7F) in the lungs of two idiopathic PAH (IPAH) patients compared to failed donor (FD) control specimens. AIP1 was highly expressed in the VSMCs of both IPAH and FD control lung tissue (Figure 7G). VSMCs lining remodeled vessels in IPAH displayed stronger AIP1 staining compared to VSMCs of normal arteries in FD controls (Figure 7G). Endothelial cell AIP1 was observed in normal, small lung vessels of FD controls but was largely absent in IPAH (Figure 7G). The increase in AIP1 expression in VSMCs and decrease in endothelial cells, while consistent with elevated AIP1 levels in SuHx rat whole lung tissue and AIP1 loss in *PHD2*-silenced human LMVECs, was notable and requires further investigation.

It has been reported that the E3-ubiquitin ligase SMURF1 (SMAD ubiquitin regulatory factor 1) targets AIP1 for proteasomal degradation.^46^ Consistent with this, silencing *PHD2* increased SMURF1 protein levels (Figure S9A), while knocking down *SMURF1* increased AIP1 protein levels in LMVECs, as was also seen for SMAD1 (Figure S9B), another known SMURF1 target protein. Moreover, *SMURF1* silencing reversed the apoptosis-resistant phenotype of *PHD2*-silenced LMVECs (Figure S9C). These results implicate SMURF1 in *PHD2* knockdown induced AIP1 loss and the pseudohypoxic, endothelial phenotype described here.

## DISCUSSION

In this study, we demonstrated that PHD2 deficiency leads to an apoptosis-resistant, hypo-proliferative, and inflammatory endothelial cell phenotype independent of HIF signaling. Mechanistically, as summarized in the schematic Figure 8D, PHD2 deficiency activated AKT and reduced AIP1 protein expression; decreased AIP1 in turn contributed to AKT and ERK activation, and JNK inactivation, which were linked to important phenotypic aspects of endothelial dysfunction including apoptosis resistance and inflammation, a hallmark feature of PAH.

Hypoxia and pseudohypoxia, both of which inactivate the PHD/VHL pathway, have been shown to alter a variety of cellular processes. In human umbilical vein endothelial cells, short exposures (6-12 h) to 5% and 10% O_2_ were found to activate anti-apoptotic mechanisms via NFkB and HIF1a activation, while severe hypoxia and longer exposures to mild hypoxia induced apoptosis, through large increases in HIF1a and accumulating DNA damage.^47^ Human PAECs exposed to hypoxia (5% and 10% O_2_) were found to proliferate over 48 h compared to normoxic cells, an effect inhibited by HIF2α knockdown.^48^ However, exposure of human dermal microvascular endothelial cells to DMOG, a cell permeable PHD2 inhibitor, blocked mtROS generation independent of HIF signaling^49^ and significantly reduced their proliferation over 48-72 h.^50^ In mouse islet endothelial cells, PHD2 silencing was shown to enhance hypoxia-induced proliferation over 72 h independent of HIF signaling.^51^ Over a similar timeframe, heterozygous PHD2 knockout reduced mouse lung endothelial cell apoptosis triggered by serum starvation and inhibited proliferation in response to VEGF.^52^ Herein, PHD2 knockdown or inhibition caused an apoptosis-resistant, hypo-proliferative, and inflammatory phenotype in human LMVECs and PAECs over 72 h, which was not reversed by blocking HIF signaling. On the contrary, *HIF2α* or *HIF1β* silencing increased apoptosis resistance. Furthermore, the specific HIF2α inhibitor, PT2567, or *HIF2α* knockdown, as reported in human blood outgrowth endothelial cells,^53^ inhibited proliferation. These observed discrepancies among different endothelial cell types and hypoxic or pseudohypoxic cellular contexts are further evidence that mechanisms and pathways other than HIFα mediate important aspects of PHD/VHL signaling. Highlighting the importance of HIF-independent effects resulting from PHD/VHL signaling, cellular transcriptomic profiles have found that many genes are differentially regulated by hypoxia or pseudohypoxia, yet are unaffected by HIF silencing.^54^ A recent study of pulmonary arteries from patients with PH found elevated HIF proteins, markedly increased cytokines/chemokines, and anti-apoptosis genes. However, HIF inhibition did not reverse the expression of these genes, or the aberrant phenotype of fibroblasts cultured *ex vivo.*^55^

As already noted here and by others,^26, 56^ PHD2 loss or its inhibition by small molecule inhibitors or hypoxia leads to HIF-independent AKT activation. PHD2 activation via its cofactor α-ketoglutarate inactivates AKT and rescues mice from lung inflammation.^56^ Consistent with these findings, *PHD2* silencing or inhibition by DMOG in LMVECs induced AKT activation and triggered an IFN-biased inflammatory response. Importantly, these effects of *PHD2* silencing in the present study were not altered by *HIF2α* siRNA silencing or PT2567, a selective HIF2α inhibitor. In a similar HIF2α-independent manner, *PHD2* silencing in LMVECs activated ERK while suppressing JNK and reducing AIP1 protein levels without altering AIP1 mRNA expression. Likewise, DMOG mimicked all these effects of *PHD2* silencing except for ERK phosphorylation, indicating that *PHD2* silencing and its inhibition by DMOG are not entirely equivalent.

As a Ras GAP scaffolding protein, AIP1 can directly interact with AKT1 and the p85 subunit of PI3K, thereby inhibiting the PI3K-AKT axis.^39^ AIP1 GAP activity further enforces this inhibition by reducing RAS-dependent activation of PI3K p100α subunit^30^ and negatively controlling ERK.^57^ Binding both ASK1 and PP2A, AIP1 facilitates removal of the inhibitory ASK1-S967 phosphorylation and accordingly activates the ASK1-JNK pathway.^40^ Loss of AIP1 also activates IFN/STAT signaling.^42^ Consistent with these previous results, *AIP1* silencing, like PHD2, indeed activated AKT and ERK, and inhibited JNK, and activated STAT1/3 in our study. Co-silencing showed even more pronounced, concordant effects, as revealed by microarray analysis. STAT1/3 activation was also detected in LMVECs isolated from APAH patients. These abnormalities were accompanied by apoptosis resistance and hypo-proliferation in LMVECs, suggesting that AIP1 loss in PHD2-deficient endothelial cells contributes to AKT, ERK, and STAT activation, JNK suppression, and subsequent cellular phenotypic aberrations.

AKT activation has been found in the vasculopathic pulmonary vessels of hypoxic mice,^58^ SuHx rats^59, 60^ and patients with IPAH.^29, 59^ Deletion of *Akt1*^58^ or pharmacological inhibition of PI3K, an upstream AKT activator, attenuates vascular remodeling and the development of PAH in rodents.^59^ Moreover, AKT and STAT1/3 activation was recently demonstrated in dermal fibroblasts from patients with HPAH due to CAV1 loss-of-function mutations.^29^ Consistent with these prior findings, the present study showed augmented activation of AKT and STAT3 in lung homogenates of SuHx rats. AKT and STAT activation in these tissues was associated with reduced PHD2 levels, supporting the link between PHD2 deficiency and increased AKT and STAT activity in PAH. Notably, the importance of PI3K/AKT signaling in STAT activation, as seen in our current study, has been reported by others. STAT1 requires phosphorylation at tyrosine 701 and serine 727 for full activation. AKT directly phosphorylates STAT1 at serine 727 ^61^ and IRF3 at serine 386.^62^ Phosphorylated IRF3 induces expression of IFNs which activates JAK with subsequent phosphorylation of STAT1 at tyrosine 701 and STAT3 at tyrosine 705.^62^ Additionally, AKT reinforces STAT signaling by promoting mRNA translation of IFN-stimulated genes via phosphorylation of mTOR/p70 S6 kinase.^63^

AIP1 is expressed in both endothelial cells and VSMCs. Mice with global deletion of *Aip1* exhibit reduced retinal angiogenesis^64^ and enhanced inflammation and endothelial dysfunction induced by hyperlipidemia.^65^ Endothelial cells isolated from *Aip1*-knockout mice or *AIP1*-silenced human endothelial cells all show attenuated proliferation,^64^ but VSMCs in endothelial cell-specific *Aip1* knockout mice are highly proliferative due to AIP1 loss-induced endothelial inflammation.^41^ In accordance with these results, human LMVECs whose AIP1 was suppressed by siRNA silencing of either *AIP1* directly or *PHD2*, displayed hypo-proliferative phenotypes. Transcriptomic profiling of these cells revealed repression of E2F and MYC target genes, which are known to promote cell proliferation. Multiple pathways may contribute to the observed repression of E2F and MYC signaling in AIP1- and PHD2-deficient LMVECs. For example, type I or type II IFNs, which were both activated in these cells, have been previously shown to inhibit E2F- and MYC-mediated gene transcription.^66, 67^

Like *PHD2*-silenced LMVECs, AIP1 protein levels assessed by immunohistochemistry were markedly decreased in the pulmonary endothelium of *Tie2-Cre*/*Phd2* knockout mice. In contrast, lung homogenates from SuHx rats, which had decreased PHD2 expression, were found to have increased levels of AIP1. This apparent paradox might be explained by the crosstalk between endothelium and its underlying vascular smooth muscle. While endothelial PHD2 deficiency suppresses AIP1 expression in these cells, the resulting inflammatory endothelium may induce VSMC hyper-proliferation and AIP1 expression. The reduced levels of PHD2 and AIP1 found in the pulmonary endothelium of patients with IPAH was associated with VSMC proliferation and strong AIP1 staining, supporting this explanation. Notably, PHD2 deficiency caused a reduction in AIP1 protein, but not mRNA in human LMVECs, suggesting post-translational regulation of AIP1 by PHD2. SMURF1, an E3-ubiquitin ligase, can target AIP1 for proteasomal degradation, which is enhanced by AKT phosphorylation of SMURF1 and AIP1.^46, 68^ Importantly, SMURF1 is increased in the tissues of patients with PAH and *Smurf1* deletion protects mice from PAH,^69^ indicating a critical role of SMURF1 and its degradation targets in disease development. Consistent with these previous findings, silencing *PHD2* increased SMURF1 protein, while knocking down *SMURF1* increased AIP1 protein in LMVECs. Moreover, knocking down *SMURF1* reversed the apoptosis-resistant phenotype of *PHD2*-silenced cells.

In summary, PHD2 deficiency-induced pseudohypoxia in human LMVECs and PAECs resulted in an apoptosis-resistant, inflammatory, and hypo-proliferative phenotype. Investigation of this *in vitro* cellular model revealed that AKT and STAT activation as well as AIP1 loss, but not HIF signaling, were critical to these phenotypic abnormalities. Two different rodent models of PAH and lung tissues from patients with PAH supported potential contributions from PHD2 loss and the activation of AKT and STAT signaling, with AIP1 dysregulation perhaps playing a more complex, cell-type specific role. Although HIF2α and/or HIF1α are widely accepted as important therapeutic targets, our current study indicates that blocking HIF signaling alone may not be sufficient to prevent or reverse pathologic vascular remodeling in PAH. Further study and therapeutic strategies targeting PHD2, AKT and AIP1 are warranted.

## SUPPLEMENTARY MATERIAL

Supplemental information can be found online.

## ACKNOWLEDGEEMNTS

The research was funded by the National Institutes of Health (NIH).

## AUTHOR CONTRIBUTIONS

S.W. and R.L.D. conceived and designed the research; S.W. wrote the manuscript; R.L.D. and J.M.E. commented and edited the manuscript; Y-Y.Z. contributed mice lung tissue slides and edited the manuscript; S.W., K.S.A., L-Y.C., M.A.H.S., G.A.F., C.L.W., J.T., Z-X.Y. and K.T. performed experiments and commented on the manuscript; S.W. and C.Y.D performed data analysis. All authors have read and agreed to the publication of the manuscript.

## DECLARATION OF INTERESTS

No conflicts of interest are declared by the authors.

## DATA AVAILABILITY STATEMENT

Data are available in the methods and/or supplementary material of this article.

## Notes

### Competing Interest Statement

The authors have declared no competing interest.

