## Supplementary Figures and Tables for "Endothelial PHD2 deficiency induces apoptosis resistance and inflammation via AKT activation and AIP1 loss independent of HIF2α"

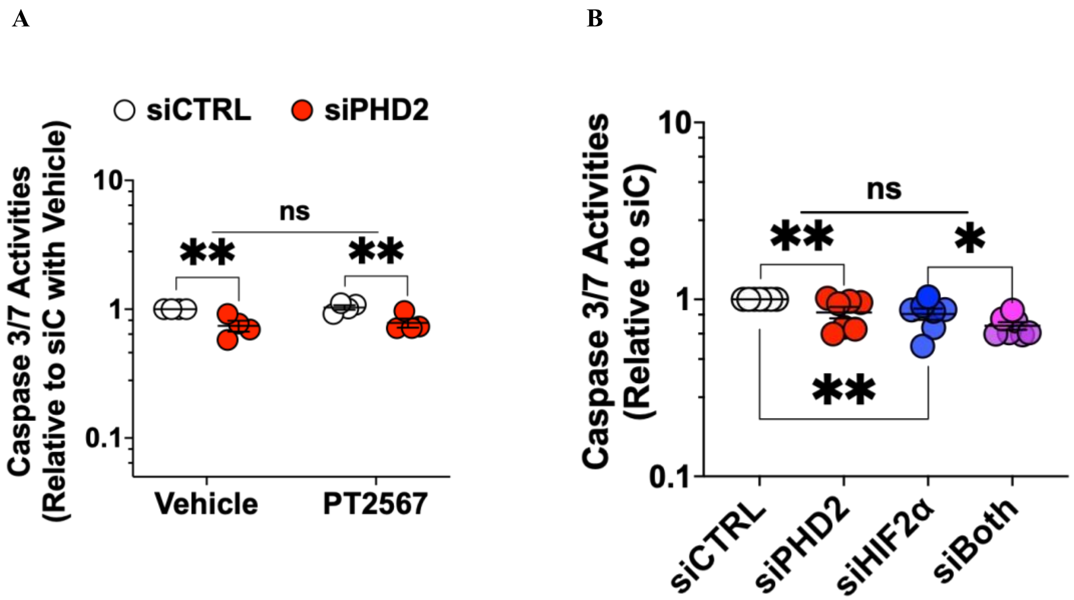


**Figure S1. HIF2α inhibition or silencing does not reverse apoptosis resistance induced by *PHD2* silencing in human pulmonary artery endothelial cells.**

Cells were transfected with scrambled control (siCTRL), *PHD2* (siPHD2), *HIF2*α (siHIF2α) siRNA, or their combination for 48 h. (A) Cells were further incubated for 24 h in serum- and growth factor-free medium (SFM-GF) with PT2567 (10 μM) or vehicle for measurement of caspase3/7 activation (n = 4). (B) Cells were further incubated in SFM-GF for 24 h for measurement of caspase3/7 activation (n = 6). Caspase activity was measured using the Caspase-Glo3/7 Assay. Data present mean ± SEM. * $p < 0.05$, ** $p \leq 0.01$. Two-way ANOVA with post-hoc t-test.

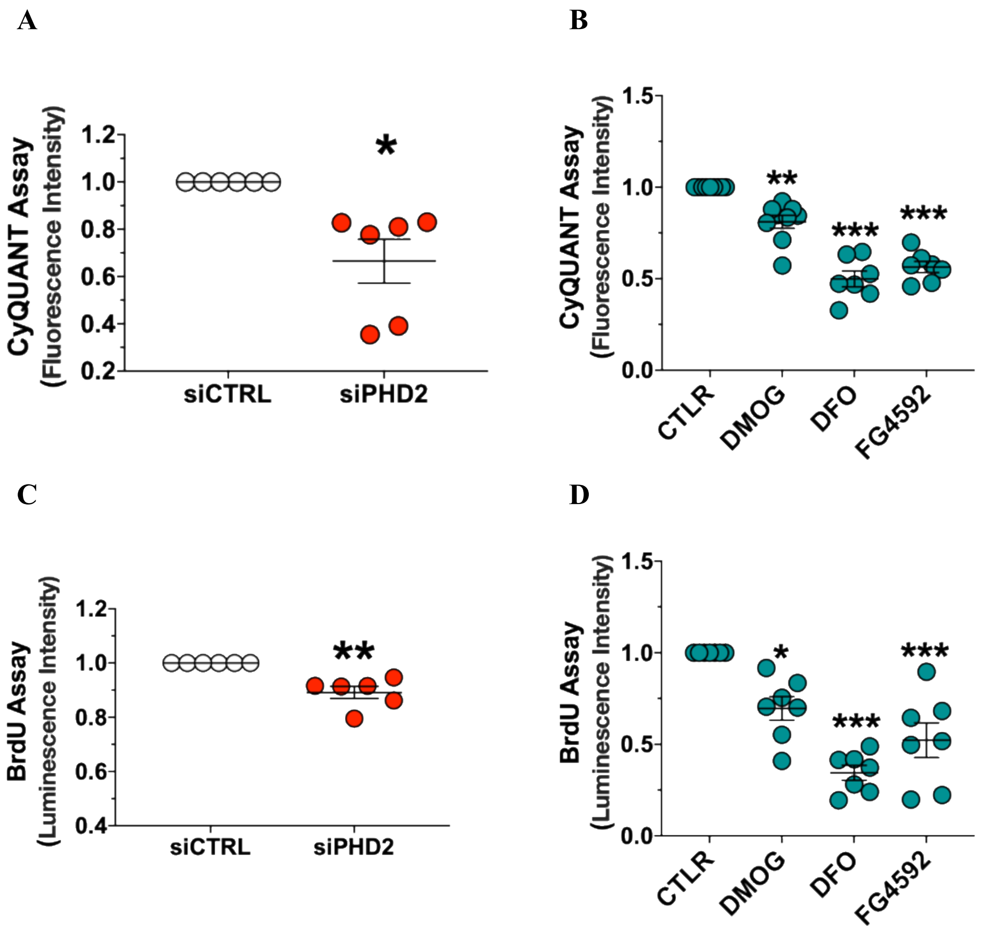


**Figure S2.** ***PHD2*** **silencing and inhibition inhibits cell proliferation in human lung microvascular endothelial cells.**
(A) Cells were transfected with scrambled control (siCTRL) or *PHD2* (siPHD2) siRNA for 72 h and cell proliferation was evaluated using the CyQUANT Assay (n = 6). (B) Cells were incubated for 72 h with different PHD2 inhibitors dimethyloxalylglycine (DMOG, 100 μM), deferoxamine (DFO, 100 μM), roxadustat (FG-4592, 50 μM) or vehicle (CTRL), and cell proliferation was evaluated as in (A) (n = 7). (C) *C*ells were transfected for 48 h, serum starved for 24 h, and labelled with BrdU for 24 h for proliferation assay (n = 6). (D) Cells were incubated for 48 h with DMOG, DFO, FG-4592 or CTRL, serum starved for 24 h, and labelled with BrdU for 24 h for proliferation assay. Data present mean ± SEM (n = 7). *$p < 0.05$, **$p \leq 0.01$, ***$p \leq 0.001$. Paired t-test (A and C), one-way ANOVA with post-hoc Dunnett's test (B and D).

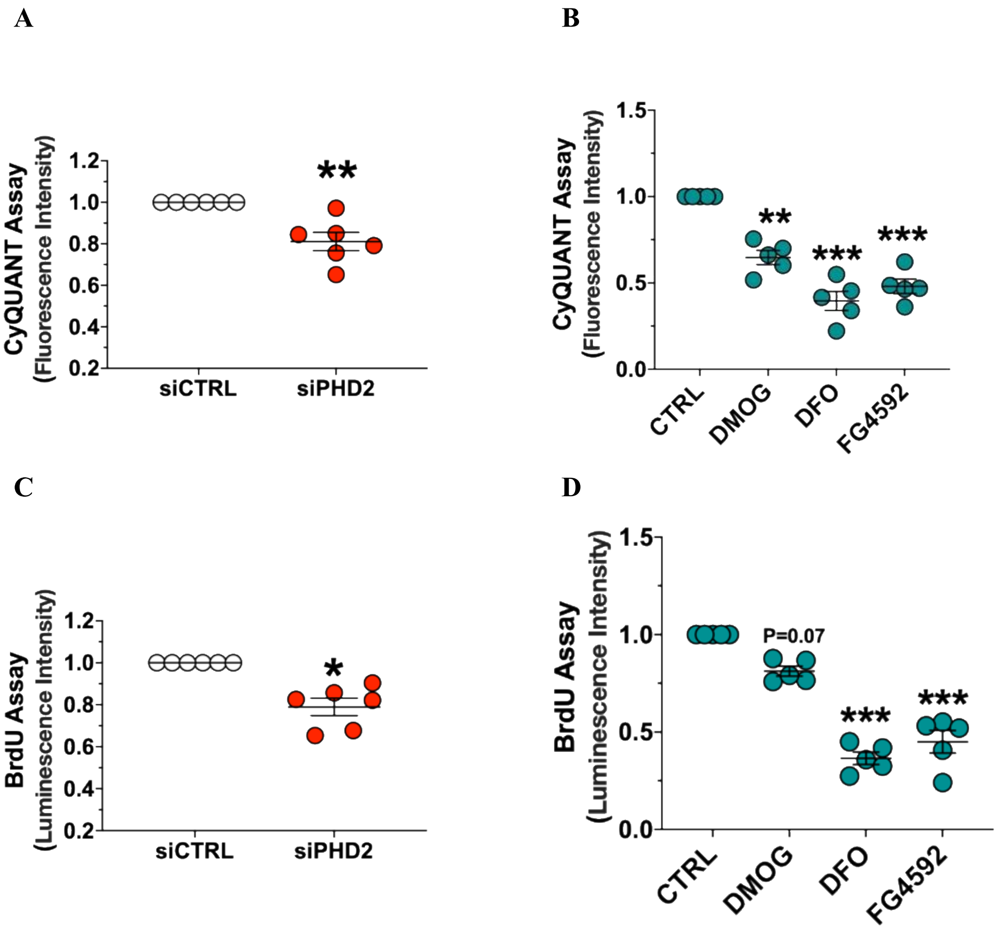


**Figure S3.** ***PHD2*** **silencing and inhibition inhibits cell proliferation in human pulmonary artery endothelial cells.**

(A) Cells were transfected with scrambled control (siCTRL) or *PHD2* (siPHD2) siRNA for 72 h and cell proliferation was evaluated using the CyQUANT Assay (n = 6). (B) Cells were incubated for 72 h with different PHD2 inhibitors dimethyloxalylglycine (DMOG, 100 μM), deferoxamine (DFO, 100 μM), roxadustat (FG-4592, 50 μM) or vehicle (CTRL), and cell proliferation was evaluated as in (A) (n = 7). (C) *C*ells were transfected for 48 h, serum starved for 24 h, and labelled with BrdU for 24 h for proliferation assay (n = 6). (D) Cells were incubated for 48 h with DMOG, DFO, FG-4592 or CTRL, serum starved for 24 h, and labelled with BrdU for 24 h for proliferation assay. Data present mean ± SEM (n = 7). *$p < 0.05$, **$p \leq 0.01$, ***$p \leq 0.001$. Paired t-test (A and C), one-way ANOVA with post-hoc Dunnett's test (B and D).

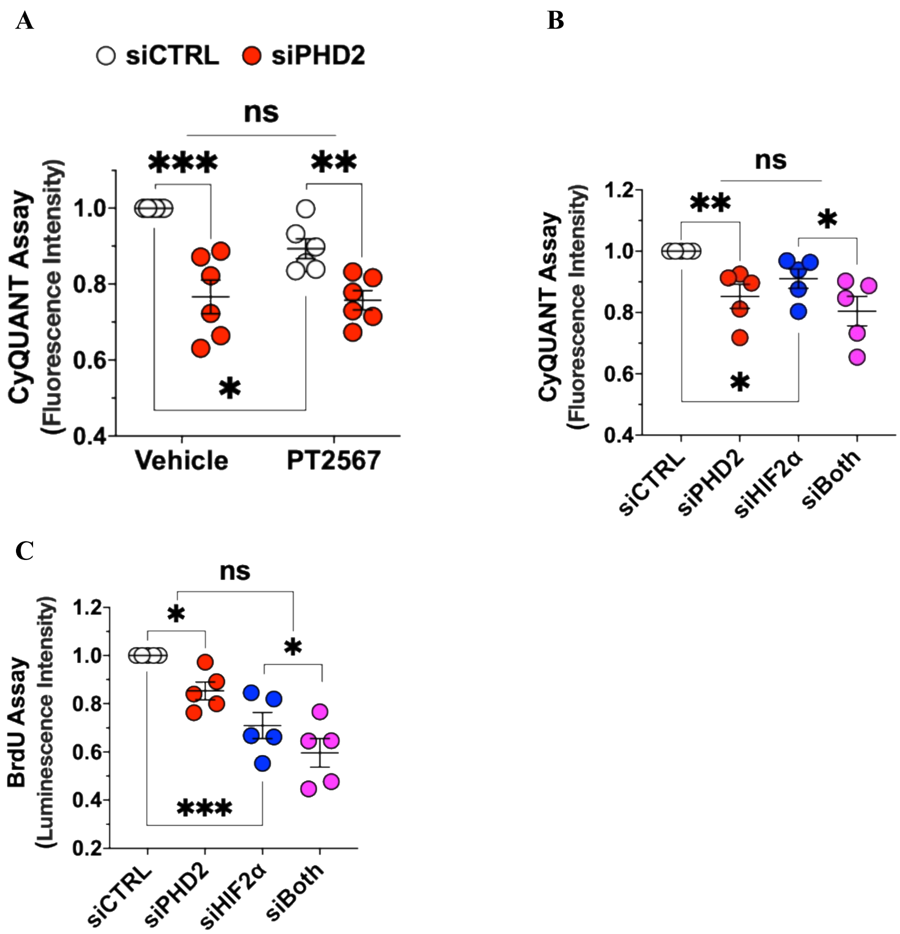


**Figure S4. HIF2α inhibition and silencing does not alter the hypo-proliferative phenotype in *PHD2*-silenced human lung microvascular endothelial cells.**

(A) Cells were transfected with scrambled control (siCTRL) or *PHD2* (siPHD2) for 72 h for proliferation assay using the CyQUANT Assay. HIF2α inhibitor PT2567 (10 μM) or control vehicle were added at 6 h post-transfection (n = 6). (B) Cells were transfected with siCTRL, siPHD2, *HIF2a* (siHIF2a) siRNA or their combination for 72 h for proliferation as in (A) (n = 5). (C) Cells were transfected with siCTRL, siPHD2, siHIF2α or their combination for 48 h, serum starved for 24 h, and labelled with BrdU for 24 h for proliferation assay (n = 5). Data present mean ± SEM. *$p < 0.05$, **$p \leq 0.01$, ***$p \leq 0.001$. Two-way ANOVA with post-hoc t-test.

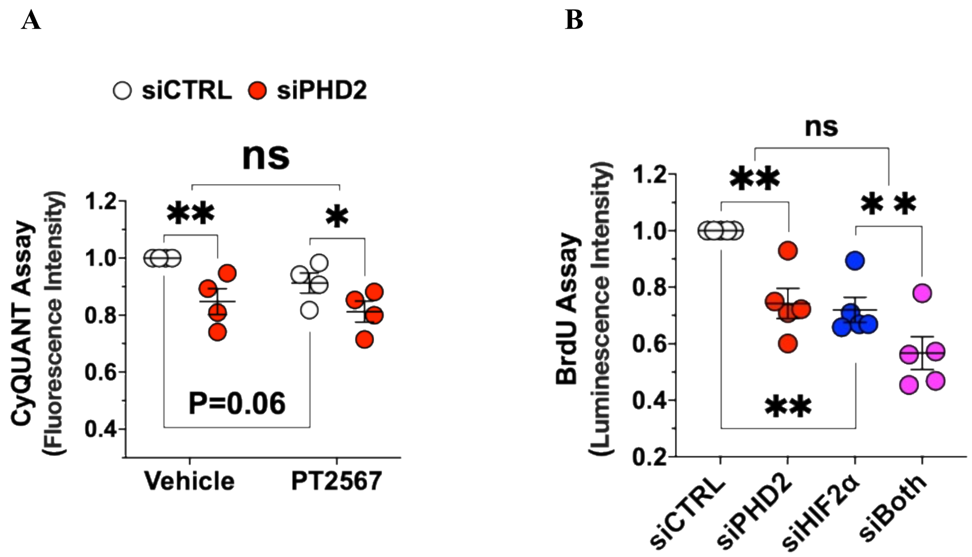


**Figure S5. HIF2α inhibition and silencing does not alter the hypo-proliferative phenotype in *PHD2*-silenced human pulmonary arterial endothelial cells.**

(A) Cells were transfected with scrambled control (siCTRL) or *PHD2* (siPHD2) for 72 h for proliferation assay using the CyQUANT Assay. HIF2α inhibitor PT2567 (10 μM) or control vehicle were added at 6 h post-transfection (n = 4). (B) Cells were transfected with siCTRL, siPHD2, *HIF2α* (siHIF2α) siRNA or their combination for 48 h, serum starved for 24 h, and labelled with BrdU for 24 h for proliferation assay (n = 5). Data present mean ± SEM. *$p < 0.05$, **$p \leq 0.01$. Two-way ANOVA with post-hoc t-test.

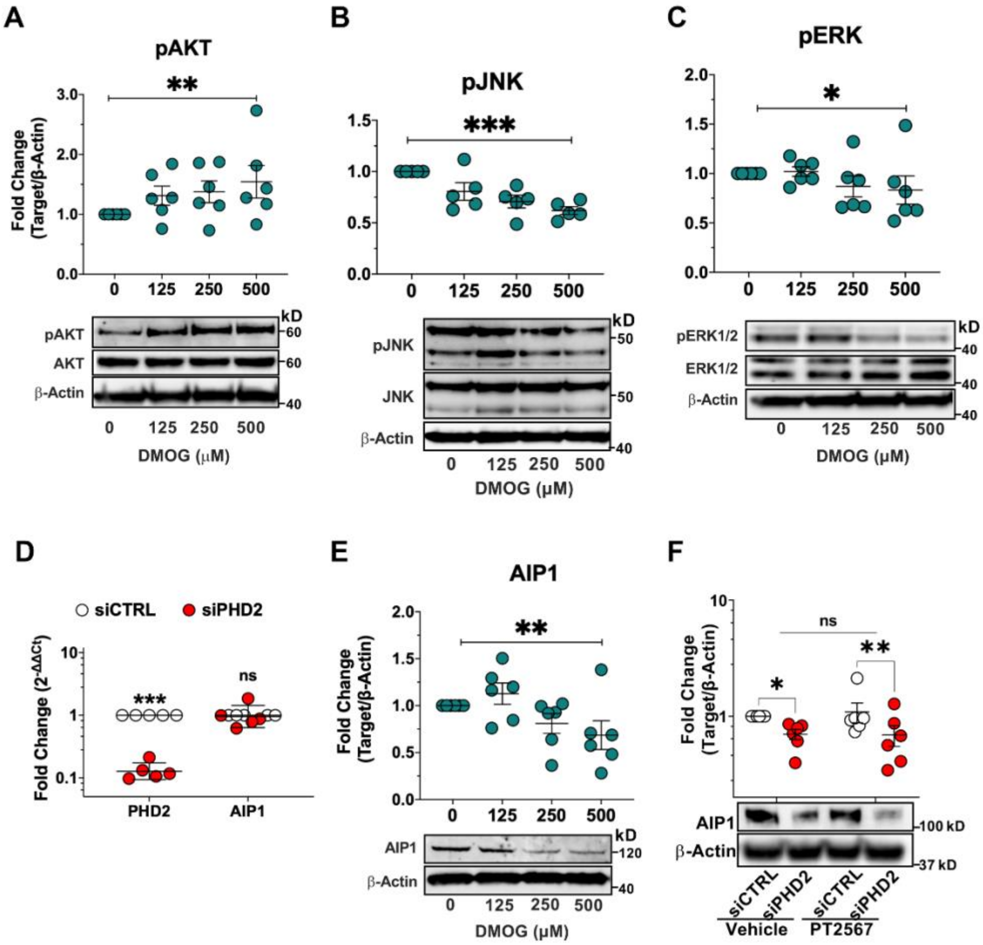


**Figure S6. The effects of PHD2 silencing and inhibitors of PHD2 or HIF2α on AKT, JNK, and ERK activation and AIP1 expression in human lung microvascular endothelial cells.**
(A-C) Cells were incubated for 24 h with different concentrations of dimethyloxalylglycine (DMOG), a PHD2 inhibitor. Activation of AKT (pAKT-S473), JNK (pJNK-T183/Y185), and ERK (ERK (pERK-T202/Y204) were assessed by Western blotting (n = 5-6). (D) *C*ells were transfected with control (siCTRL) or *PHD2* (siPHD2) siRNA for 48 h. *PHD2* and *AIP1* mRNAs are measured by qPCR and presented as the geometric mean ± geometric SD (n = 5). (E) Cells were incubated and treated as in (A), and AIP1 protein levels were assessed by Western blotting (n = 6). (F) Cells were transfected with siCTRL or siPHD2 for 48 h and incubated for another 24 h with HIF2α inhibitor PT2567 (10 μM) or vehicle to measure AIP1 protein by Western blotting (n = 6). Western blots protein levels were normalized to β-actin and presented as mean ± SEM. *$p < 0.05$, **$p \leq 0.01$, ***$p \leq 0.001$. Linear regression (A, B, C, and E), paired t-test (D), two-way ANOVA with post-hoc t-test (F).

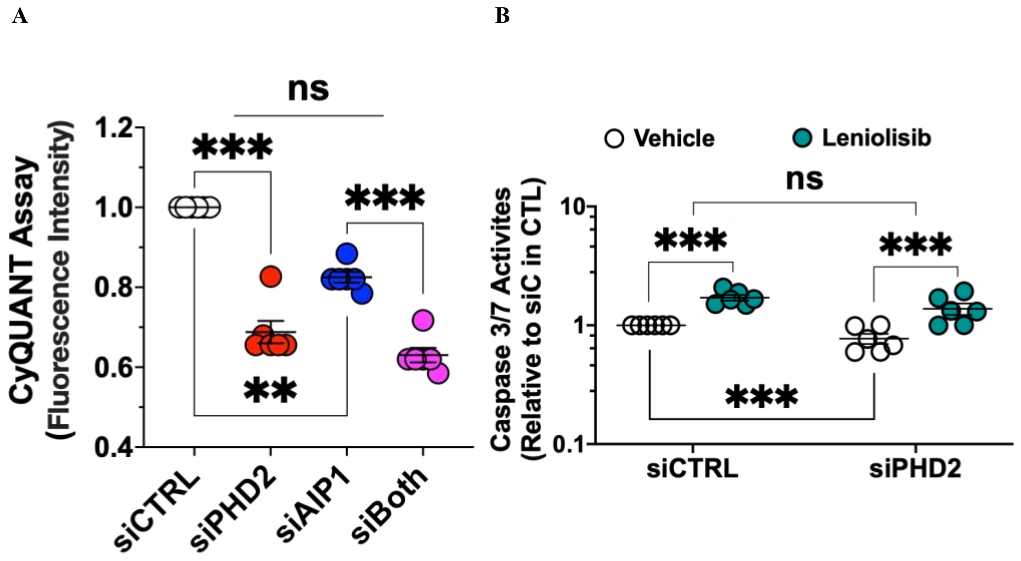


**Figure S7. The effects of *PHD2* or *AIP1* silencing and AKT inhibition on proliferation and apoptosis in human lung microvascular endothelial cells.** (A) Cells were transfected with scrambled control (siCTRL), *PHD2* (siPHD2), *AIP1* (siAIP1), or their combination for 72 h, and proliferation was evaluated using the CyQUANT Assay (n=5). (B) Cells were transfected with siCTRL or siPHD2 for 48 h and then incubated for another 24 h in serum- and growth factor-free medium with control vehicle or leniolisib (5 μM), an inhibitor of AKT activator PI3Kδ. Caspase activation was then measured using the Caspase-Glo3/7 Assay (n=6). Data present mean ± SEM. $^{**}p \leq 0.01$, $^{***}p \leq 0.001$. Two-way ANOVA with post-hoc t-test.

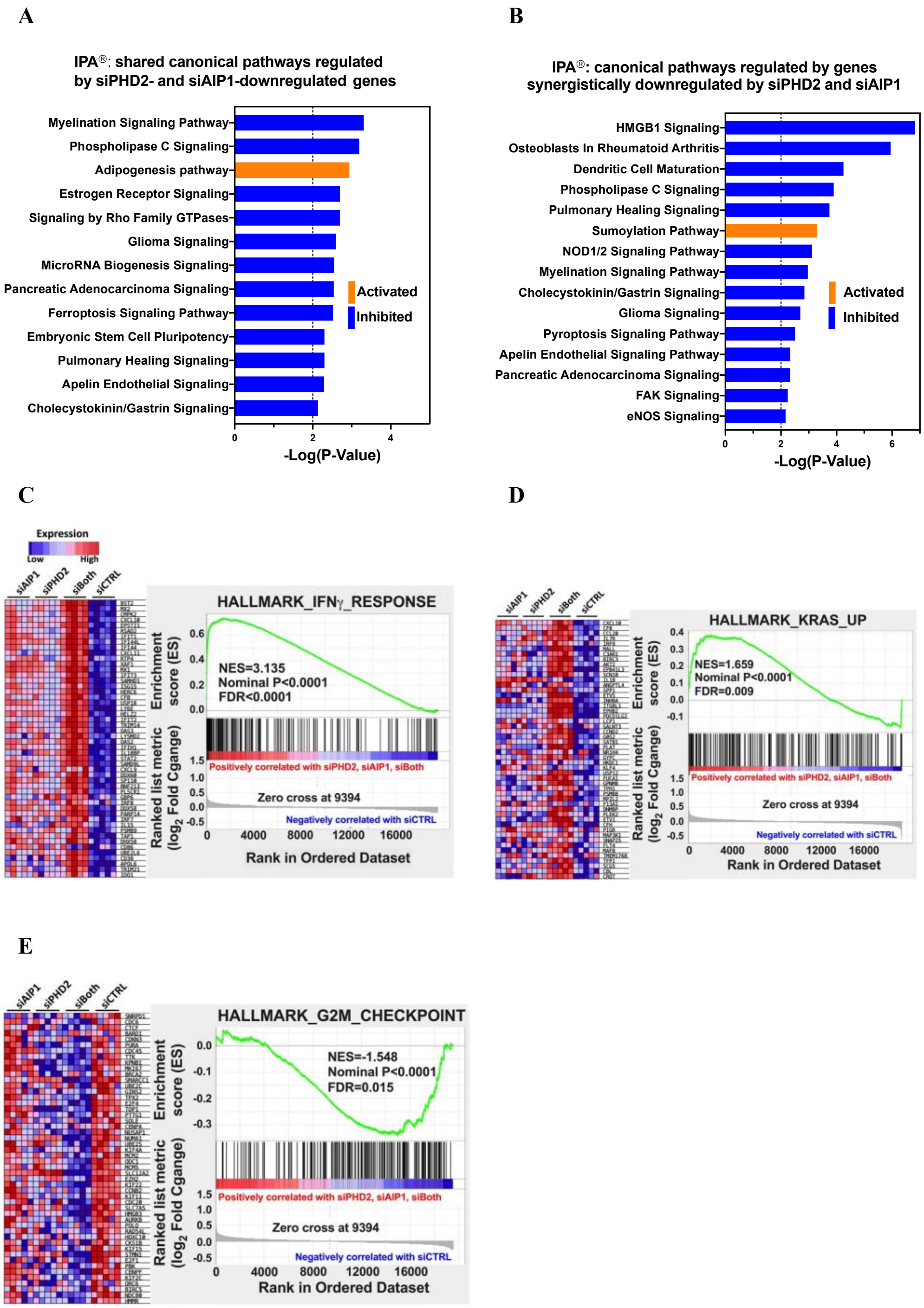


**Figure S8. Transcriptome analysis of human lung microvascular endothelial cells silenced for *PHD2*, *AIP1*, and both using microarrays.**

(A, B) Top canonical pathways enriched by Ingenuity pathway analysis (IPA, $p \leq 0.01$) for genes downregulated by either *PHD2* (siPHD2) or *AIP1*(siAIP1) silencing (A) and genes synergistically downregulated by *PHD2* and *AIP1* silencing (B). (C-E) Gene set enrichment analysis (GSEA, siPHD2, siAIP1, and siBoth vs. siRNA siCTRL). Expression heat maps of the top 50 marker genes and enrichment plots for the enriched pathways displayed significant activation of IFNγ (C) and KRAS (D), and suppression of G2M checkpoint (E) by silencing of *PHD2*, *AIP1*, and both.

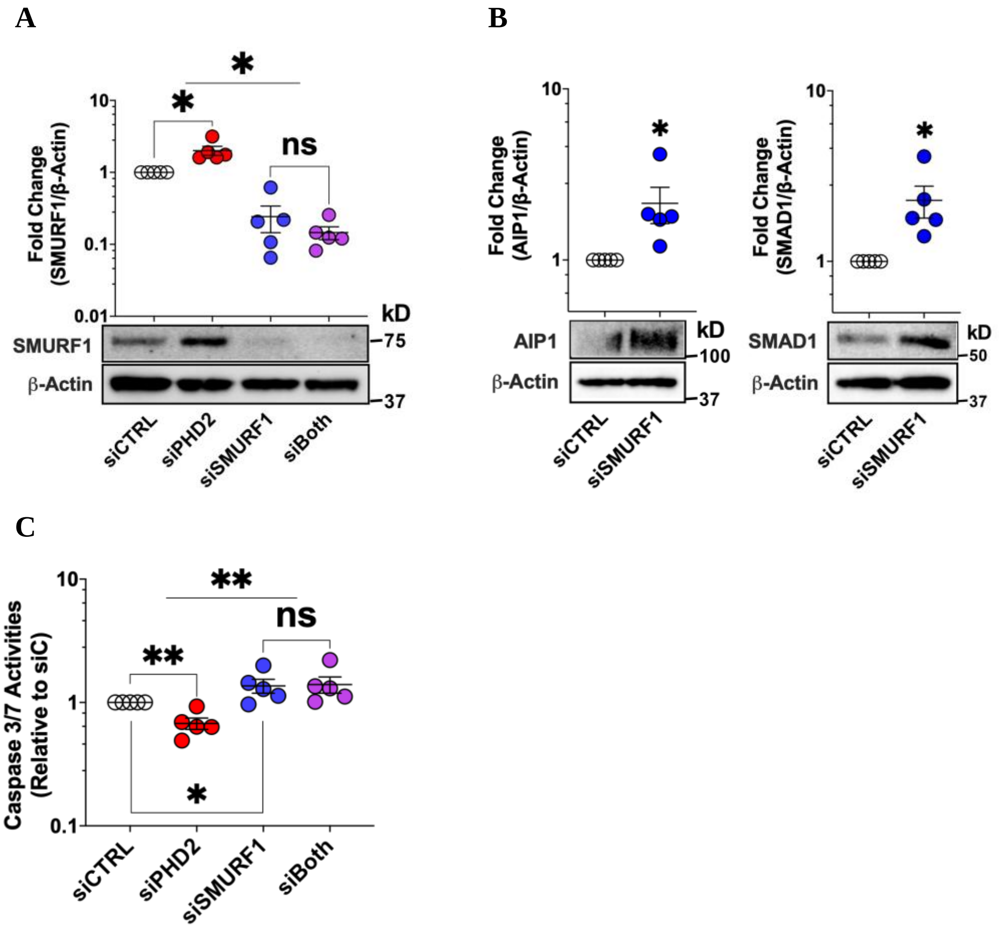


**Figure S9. *PHD2* silencing increased SMURF1, while *SMURF1* knockdown increased AIP1 and reversed *PHD2* silencing-induced apoptosis resistance in human lung microvascular endothelial cells.** Cells were transfected with scrambled control (siCTRL), *PHD2* (siPHD2), *SMURF1* (siSMURF1) siRNA or both for 48 h. (A, B) Whole cell lysates were collected for Western blotting of SMURF1, AIP1, and SMAD1. Densitometric quantifications of protein expression were normalized to b-actin and representative Western blots shown. (C) Cells were further incubated for 24 h in serum- and growth factor-free medium to measure caspase activities using the Caspase-Glo3/7 Assay. Data present mean ± SEM. *p < 0.05, **p ≤ 0.01 (n = 5 for all). Two-way ANOVA with post-hoc t-test (A and C)**,** paired t-test (B).

**Supplementary Tables:**

**Table S1. Key resource table**

| REAGENT or RESOURCE | SOURCE | IDENTIFIER |
|---|---|---|
| Antibodies | | |
| PHD2/EGLN1 | Cell Signaling | Cat#4835; RRID: AB_10561316 |
| PHD2/EGLN1 (IF) | LSBio | Cat#LS-C331303; RRID : AB_3076256 |
| HIF2$\alpha$/EPAS1 | Abcam | Cat#ab199; RRID: AB_302739 |
| GLUT1/SLC2A1 | Cell Signaling | Cat#12939; RRID: AB_2687899 |
| HK2 | Cell Signaling | Cat#2867; RRID: AB_2232946 |
| PKM2 | Cell Signaling | Cat#4053; RRID: 1904096 |
| LDHA | Cell Signaling | Cat#2012; RRID: AB_2137173 |
| AIP1/DAB2IP | Abcam | Cat#ab87811; RRID: AB_2041032 |
| AIP1/DAB2IP (IF) | Proteintech | Cat#23582-1-AP ; RRID : AB_2879300 |
| $\beta$-Actin | Sigma | Cat#A3854; RRID: AB_262011 |
| CXCR4 | Novus Biologicals | Cat#NBP1-77067; RRID: AB_11005253 |
| HIF1$\beta$ /ARNT | Cell Signaling | Cat#5537; RRID: AB_10694232 |
| AKT, total | Cell Signaling | Cat#9272; RRID: 329827 |
| Phospho-AKT (Ser473) | Cell Signaling | Cat#9271; RRID: AB_329825 |
| JNK, total | Cell Signaling | Cat#9252; RRID: AB_2250373 |
| JNK1, total | Cell Signaling | Cat#3708; RRID: AB_1904132 |
| Phospho-JNK (Thr183/Tyr185) | Cell Signaling | Cat#4668; RRID: AB_823588 |
| p44/42 MAPK (ERK1/2) | Cell Signaling | Cat#9102; RRID: AB_330744 |
| Phospho-p44/42 (Thr202/Tyr204) | Cell Signaling | Cat#9101; RRID: AB_331646 |
| Phospho-STAT1 (Tyr701) | Cell Signaling | Cat#7649; RRID: AB_10950970 |
| Total STAT1 | Cell Signaling | Cat#9172; RRID: AB_2198300 |
| Phospho-STAT3 (Tyr705) | Cell Signaling | Cat#9145; RRID: AB_2491009 |
| Total STAT3 | Cell Signaling | Cat#9139; RRID: AB_331757 |
| SMURF1 | Abcam | Cat#ab57573; RRID: AB_945548 |
| SMAD1 | Cell Signaling | Cat#9743; RRID: AB_2107780 |
| von Willebrand Factor (vWF) | BIO-RAD | Cat# AHP062; RRID: AB_322241 |
| Donkey anti-sheep | Abcam | Cat#ab150177; RRID: AB_2801320 |
| Donkey anti-rabbit | Abcam | Cat# ab150076; RRID: AB_2782993 |
| Biological samples | | |
| Lung sections of IPAH patients | Pulmonary Hypertension Breakthrough Initiative (PHBI) Penn Cell Center, | PHBI#: BA023, ST052, UA026, UC007, VA015 |
| Lung sections of failed donor controls | PHBI | PHBI#: AH028, BA062, UA020, UC011, VA008 |

| | | |
|---|---|---|
| Lung microvascular endothelial cells from APAH patients and failed donor controls | PHBI | Cat#02544 |
| Chemicals | | |
| SU5416 | Tocris | Cat#3037 |
| PT2567 | Peloton Therapeutics | Custom |
| Dimethyloxalylglycine (DMOG) | Cayman Chemical | Cat#71210 |
| Deferoxamine (DFO) | Cayman Chemical | Cat#14595 |
| Roxadustat (FG-4592) | Cayman Chemical | Cat#15294 |
| Leniolisib | Novartis | Custom |
| Critical commercial assays | | |
| Caspase-Glo 3/7 | Promega | Cat#G8093 |
| FITC Annexin V Apoptosis Detection Kit | BD Pharmingen | Cat#556547 |
| CyQUANT® Cell Proliferation Assay | ThermoFisher Scientific | Cat#C7026 |
| Cell Proliferation ELISA, BrdU | MilliporeSigma | Cat#11669915001 |
| CXCL10 ELISA | R&D Systems | Cat#DIP100 |
| MitoTracker Green FM | ThermoFisher Scientific | Cat#M7514 |
| MitoSOX Red | ThermoFisher Scientific | Cat#M36007 |
| DharmaFECT1 | Dharmacon | Cat#T2001-02 |
| Experimental models: Cell lines | | |
| Human lung microvascular endothelial cells (LMVECs) | Lonza | CC-2527 |
| Human pulmonary artery endothelial cells (PAECs) | Lonza | CC-2530 |
| Experimental models: Organisms/strains | | |
| Rat: Sprague-Dawley (SD), controls for PAH rat models | Charles River Laboratories | 400 |
| SU5416/hypoxia SD rat PAH models (SuHx) | This work | N/A |
| *Egln1*$^{Tie2}$ (*Phd2* CKO) mice and wild type controls | (Dai et al., 2016)[1] | N/A |
| Oligonucleotides | | |
| TaqMan primers/probes for qPCR, See Table S1 | ThermoFisher Scientific | N/A |
| siRNAs for gene silencing, See Table S2 | Dharmacon | N/A |
| Human Clariom™ S Assay | ThermoFisher Scientific | 902927 |

**Table S2. TaqMan™ Expression Assays for qPCR (Thermo Fisher Scientific)**

| Gene Symbol | Gene Name | Unique Assay ID |
|---|---|---|
| *ACTB* | β-Actin | Hs99999903_m1 |
| *B2M* | β-2-microglobulin | Hs00187842_m1 |
| *RPL13* | Ribosomal protein L13 | Hs00744303_s1 |
| *AIP1 (DAB2IP)* | ASK1-interacting protein 1 (AIP1); DAB2 interacting protein | Hs00368995_m1 |

| | | |
|---|---|---|
| *EGLN1* | Egl-9 Family Hypoxia Inducible Factor 1; Prolyl hydroxylase domain 2 (PHD2) | Hs00254392_m1 |
| *EPAS1* | Endothelial PAS domain protein 1; Hypoxia-inducible factor 2-alpha (HIF-2α) | Hs01026149_m1 |
| *HK2* | Hexokinase 2 | Hs00606086_m1 |
| *LDHA* | Lactate dehydrogenase A | Hs01378790_g1 |
| *PKM2* | Pyruvate kinase M2 | Hs00761782_s1 |
| *SLC2A1* | Solute carrier family 2 member 1; Glucose transporter 1 (GLUT1) | Hs00892681_m1 |

**Table S3. siRNA for gene silencing (Dharmacon)**

| **Gene Symbol** | **Gene Name** | **Cataglog #** |
|---|---|---|
| *PHD2 (EGLN1)* | Dharmacon | L-004276-00-0005 |
| *AIP1 (DAB2IP)* | | |
| *HIF2A (EPAS1)* | Dharmacon | L-004814-00-0005 |
| *HIF1β (ARNT)* | Dharmacon | L-007207-00-0005 |
| *SMURF1* | Dharmacon | L-007191-00-0005 |
| ON-TARGET plus Non-targeting pool | Dharmacon | D-001810-10-05 |


1. Dai, Z., et al., *Prolyl-4 Hydroxylase 2 (PHD2) Deficiency in Endothelial Cells and Hematopoietic Cells Induces Obliterative Vascular Remodeling and Severe Pulmonary Arterial Hypertension in Mice and Humans Through Hypoxia-Inducible Factor-2alpha.* Circulation, 2016. **133**(24): p. 2447-58.
